# Tox21mer, A transformer foundation model for Tox21 high-throughput concentration–response curves data

**DOI:** 10.64898/2026.06.15.732308

**Authors:** Leping Li, Jisoo Hwang, Keith Shockley, Yuanyuan Li, Alison Motsinger-Reif, Jui-Hua Hsieh, Scott Auerbach, David Reif

## Abstract

The U.S. Tox21 collaboration has generated a large reference library of high-throughput concentration–response assays. Here we present Tox21mer, a 43.5-million-parameter transformer that encodes each Tox21 concentration–response curve together with assay metadata into a 768-dimensional representation. Tox21mer was pretrained on ~2.5 million curves from 102 assay protocols and 6,727 compounds using masked-response reconstruction as the primary objective, with low-weight auxiliary supervision on assay outcome and AC_50_. To evaluate the learned representation, we trained lightweight probes on frozen embeddings from concentration–response curves of held-out compounds. The representation supported a macro-F1 of 0.985 for three-class outcome prediction (agonist, antagonist, inactive), a binary F1 of 0.994 for active/inactive prediction, and an R^2^ of 0.87 for log_10_(AC_50_). The learned embeddings formed coherent groupings by curve-class category. A masked-only pretraining variant retained near-baseline probe performance, indicating that the representation is learned largely from the self-supervised objective rather than from auxiliary labels. Ablation analyses further showed that predictive performance depends mainly on curve-level response-value distributions conditioned on assay context, with limited reliance on detailed within-curve ordering. Tox21mer thus provides a reusable foundation representation for Tox21 concentration-response data that can support extrapolation to untested compounds through integration with chemical features or distillation into chemistry-only student models for large-scale external screening.

## Introduction

The U.S. Tox21 collaboration is a joint effort from the U.S. Environmental Protection Agency (EPA), the National Institute of Environmental Health Sciences (NIEHS), the Food and Drug Administration (FDA), and the National Center for Advancing Translational Sciences (NCATS) ^1–11^. The primary goal of the Tox21 program is to develop more efficient, rapid, and human-relevant methods for predicting the health effects of chemicals, while reducing reliance on traditional animal-based models and improving protection against potentially harmful environmental substances and medicines.

The Tox21 program uses a large-scale automated robotic platform to evaluate thousands of chemicals for potential biological activity and toxicity through quantitative high-throughput screening (qHTS) ^12^. In practice, this ongoing effort has generated a standardized reference dataset of concentration–response profiles for a broad chemical library that includes industrial and environmental chemicals, pesticides, pharmaceuticals, and other compounds tested across more than one hundred toxicity-relevant biological targets and pathways ^5^. Most Tox21 assays are conducted in 1,536-well plate format and rely on reporter-gene cell lines, cell-viability assays, and other mechanistically informative cell-based readouts to detect pathway perturbation, identify bioactive compounds, and support chemical prioritization for follow-up testing and mechanistic interpretation ^1,5,13^. A typical qHTS experiment measures each compound across a 15-point concentration series within a single assay protocol, producing a concentration–response curve rather than a single-point activity call. In addition to the raw response measurements, each record is accompanied by structured assay metadata, such as protocol identity, target class, cell type, assay mode, detection technology, and ratio-flag status, and by curve-fit summaries including an activity outcome label and an AC_50_ (Activity Concentration at 50% of maximum activity) estimate. This rich data allows facilitating cross-assay comparisons, hit prioritization, and downstream computational modeling.

The breadth, heterogeneity, and incomplete coverage of the Tox21 assay compendium motivate the use of transformer-based foundation models, which can learn generalizable representations from large collections of concentration–response experiments rather than from individual assays in isolation. In this setting, the model encodes the full concentration–response vector together with assay metadata to produce a contextual per-curve CLS (class) embedding, which can subsequently be combined with chemical structure features-derived, for example, from ChemBERTa ^14^, in a downstream compound-level predictor. Such an approach has the potential to leverage shared biological structure across assays, improve robustness in data-sparse settings, and provide transferable representations for prediction, prioritization, and mechanistic inference.

Prior machine-learning studies involving Tox21 have focused primarily on compound-level toxicity prediction from chemical structure or assay activity labels ^15–28^, rather than pretraining a transformer on full qHTS concentration–response curves together with assay metadata to learn reusable curve-level representations. Here, we pretrained Tox21mer, a transformer model for Tox21 assay data, on approximately 2.5 million concentration–response curves and evaluated its performance on downstream branch-probe tasks. We treat the assay transformer as a foundation-style model: a single pretrained encoder shared across Tox21 assays, in the spirit of multitask representation learning ^29^ and modern foundation-model pretraining ^30^. We further examined the model’s learned representation through targeted comparisons that assessed the contributions of assay metadata, concentration–response information, and label structure, and contrasted the transformer with simpler multilayer perceptron baselines trained on either summary features or the raw 15-point response vector.

## Methods

### Dataset

Individual assay-protocol datasets were downloaded from the Tox21 Public Data portal (https://tripod.nih.gov/pubdata/). A total of 94 datasets were initially retrieved. Three reference datasets designed to detect assay interference (tox21-spec-hek293-p1, tox21-spec-hepg2-p1, and tox21-luc-biochem-p1) were excluded from further analysis. Three multimodal and/or multi timepoint datasets - DT40 and two real-time viability assays were then decomposed into single-modality datasets. The Tox21 DT40 assay comprises three modalities corresponding to distinct DNA repair capacities: wild-type DT40 cells, Rad54^−/−^ cells deficient in homologous recombination, and Ku70^−/−^ cells deficient in non-homologous end joining. We therefore split the DT40 dataset into three separate datasets according to sample data type: 100 series for wild-type DT40, 653 series for Rad54^−/−^, and 657 series for Ku70^−/−^. Two cell-viability assays, one performed in HEK293 cells and the other in HepG2 cells, were further divided according to the combination of assay technology (fluorescence or luminescence) and treatment duration (8, 16, or 24 h); other time points were not included. The metadata used in the analysis is provided in Supplementary **Table S1**.

We applied a series of structural, data-quality, and assay-content filters to the original 7,947,446 Tox21 concentration–response curves. The preprocessing strategy was designed to create a coherent corpus for representation learning while reducing ambiguity in the biological interpretation of each curve. Because the original Tox21 files include primary readouts, secondary/counter-screen readouts, raw channel measurements, multi-timepoint assays, and curves with uncertain activity interpretation, we restricted the foundation-model corpus to response profiles that could be consistently represented as protocol-specific concentration–response curves with interpretable assay context. First, we removed curves associated with very small structures (10,506 with fewer than three heavy atoms), ambiguous concentration annotations [3,073 lacking a curve_class2 (cc2) label or with cc2 =5], and low-purity samples [2,026,038 with purity other than A (≥ 90%) or B (~80-90%)]. We then filtered on assay content encoded in SAMPLE_DATA_TYPE, excluding raw channel readouts (2,770,160), complementary viability readouts (1,748,332), and FITC- or rhodamine-labeled channels (35,328 each). These endpoints were excluded here because they serve secondary roles in the Tox21 assay framework: the raw channel readouts (for example, ch1 and ch2) underlie the primary ratio readout, other readouts such as viability measurements are used as counterscreens that do not serve as the primary mechanistic agonist/antagonist endpoints. After preprocessing, 102 datasets were assembled into a single combined resource containing 2,499,594 assay curves (**Table 1**). The integrated dataset included protocol name, target category, cell type, assay mode, technology, sample ID, sample data ID, sample data type, assay outcome, cc2, AC_50_, efficacy, the 15 concentration–response values, CAS number, sample name, SMILES string, and Tox21 ID.

**Table 1.** Dataset Summary.

| Property | Value |
| --- | --- |
| Assay rows (raw) | 2,499,594 |
| Assay rows (after QC) | 2,489,975 |
| Unique compounds (SMILES) | 6,727 |
| Unique protocols | 102 |
| Agonist curves | 82,344 |
| Antagonist curves | 73,237 |
| Cytotoxic curves | 68,574 |
| Inactive curves | 1,052,461 |
| Unknown / inconclusive curves | 1,213,359 |
| Train / Val / Test chemical split | 4,708 / 1,009 / 1,010 |
| Train / Val / Test row counts | 1,736,846 / 355,593 / 410,158 |

After quality filtering, which required at least 12 valid response measurements per curve and agreement between cc2 and the reported assay outcome, 2,489,975 assay rows remained. Specifically, 996 rows with agonist + cc2 < 0 and 1,023 rows with antagonist + cc2 > 0 were removed. For all curves with cc2 = 4 (flat curve), we assigned the outcome as “inactive” and set AC_50_ to NaN, irrespective of whether these values were present in the original records. The above curve-level data were used to train the assay transformer.

For probe evaluation only, the eight original assay outcome strings recorded in Tox21 records ^28^ are mapped to five collapsed classes (Supplementary **Table S2**); only the three active class strings (Agonist, Antagonist, Inactive) supply supervised outcome labels during pretraining. Note that the Antagonist label in agonist mode assays show inhibition but may reflect increased cytotoxicity or promiscuous report gene inhibition rather than true antagonism. Similarly, agonist outcomes in antagonist mode assays show activation that might not represent true agonism ^28^. The cytotoxic and inconclusive classes are excluded from outcome and AC_50_ supervision to base learning on high quality signals not affected by potential cytotoxicity, but their rows are retained in the masked-response reconstruction objective. The strings for “active agonist” and “inconclusive agonist” (and “active antagonist” and “inconclusive antagonist”) are merged because the inconclusive subset includes well-characterized partial responders whose biological direction is unambiguous; collapsing them increases supervision support for the minority agonist and antagonist classes without introducing label ambiguity at the receptor-mode level.

In the probe setting only, the 102 assays were partitioned into mechanistic and non-mechanistic branches according to assay mode (Supplementary **Table S1**). This branch-level formulation was designed to enable assay-mode-specific prediction consistent with the biological interpretation of the underlying readout. Mechanistic assays were modeled as a three-class classification task with labels agonist, antagonist, and inactive, whereas non-mechanistic assays were treated as a binary classification task with labels active and inactive.

For the mechanistic probe, we further standardized supervision by retaining only robust curves: agonist and antagonist examples were restricted to cc2 values of −1.1, −1.2, −2.1, −2.2, 1.1, 1.2, 2.1, or 2.2, whereas inactive examples were restricted to cc2 = 4. We refer to this filter as `cc2 strict`. For the non-mechanistic branch probe, the original outcome labels were not used, because agonist/antagonist distinctions are not meaningful for these assays. Instead, probe labels were assigned directly from cc2: curves with `cc2=strict` were labeled active, and curves with `cc2=4` were labeled inactive. These label transformations were used only for probe evaluation and were not part of the masked-response pretraining objective. We also defined a broader `cc2 loose` set that included all valid cc2 classes (1.1, 1.2, 1.3, 1.4, 2.1, 2.2, 2.3, 2.4, 3.0, 4.0, −1.1, −1.2, −1.3, −1.4, −2.1, −2.2, −2.3, −2.4, and −3.0). Importantly, these exclusions and cc2-based filters were applied only during probing and were not used in constructing the foundation model.

### Data normalization and splitting

We implemented protocol-specific normalization with winsorization to handle outliers. For each protocol, response values were clipped to the 1^st^ and the 99^th^ percentiles computed from the training set, then standardized using protocol-specific mean and standard deviation. Concentration values were log_10_-transformed.

To prevent information leakage from chemical-structure overlap, we performed compound-level splitting by unique SMILES, so that all replicates and all assays for a given compound were assigned to the same split. The data were partitioned into training (70%), validation (15%), and test (15%) sets with a fixed random seed. Because each compound appears across multiple protocols with replicates, this yielded 4,708 / 1,009 / 1,010 compounds and 1,736,846 / 358,146 / 394,983 curve-level rows.

### Tox21mer Overview

Each Tox21 concentration-response curve enters the Tox21mer encoder as 16 tokens, a learnable [CLS] token plus 15 concentration–response (DR) tokens that combine assay-context embeddings (six categorical metadata fields), log-concentration, and z-scored response. The encoder is pretrained on the 70% training-split compounds (1,736,846 curves drawn from 6,727 SMILES) with masked-response MSE as the primary objective (20% of valid response positions per curve, reconstruct true values). Three low-weight auxiliary heads (λ = 0.1) outcome from CLS, outcome from mean-pooled DR tokens, and AC_50_ from mean-pooled DR tokens, provide additional gradient signal during pretraining (**Figure 1**). In stage 2 (probe evaluation), with backbone weights frozen, three linear probe heads, a 3-class mechanism head, a 2-class non-mechanism head, and a regression AC_50_ head, are trained from scratch on training-split CLS embeddings (5 epochs each) and evaluated on held-out test-split compounds. In stage 3, full-data run used the same architecture, optimizer, loss configuration, batch size and 50-epoch training schedule as the split-based Phase 1 pretraining were used to construct the foundation model.

**Figure 1.**
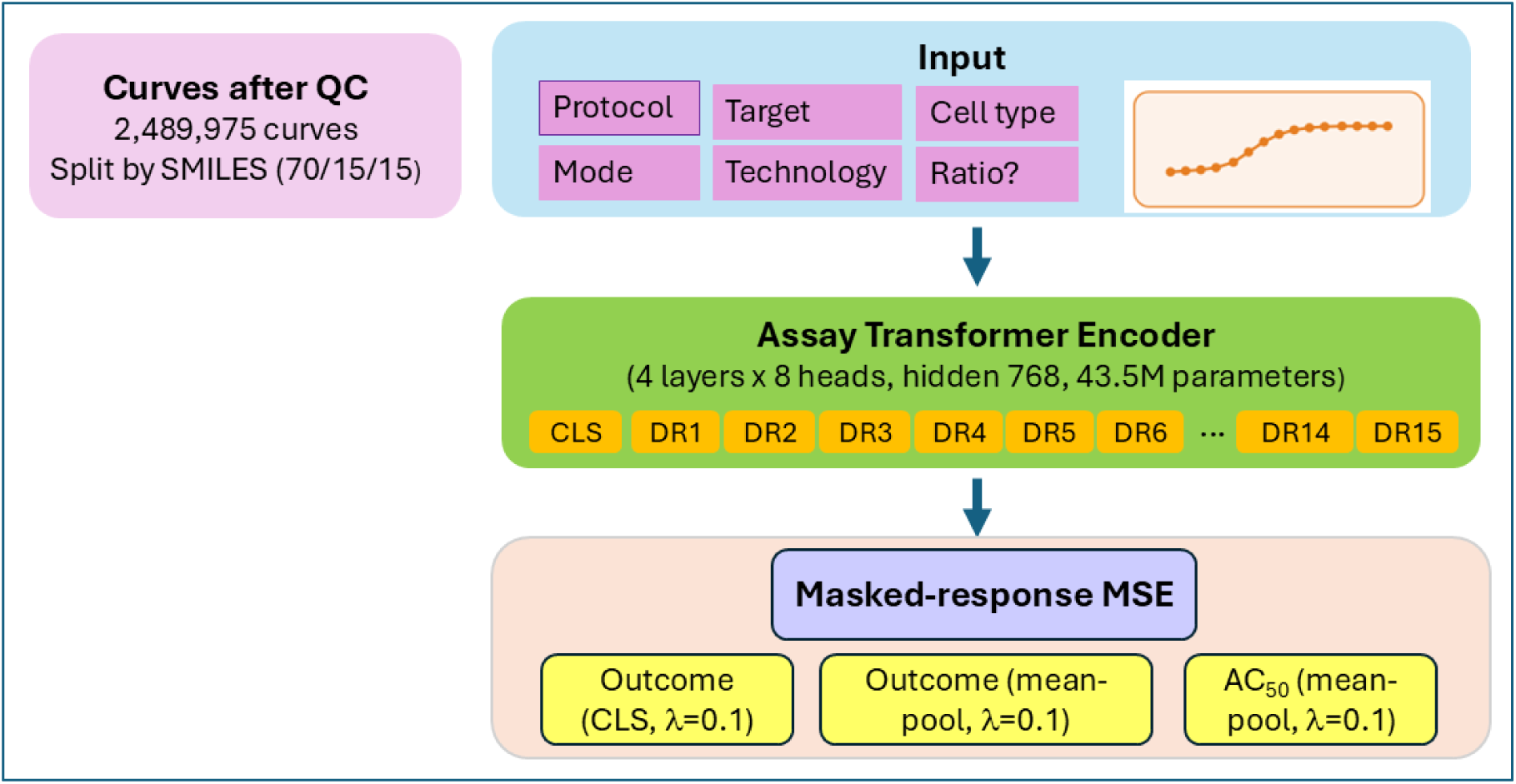
Overview of Tox21mer. Each Tox21 curve is represented as 16 tokens: one learnable [CLS] token and 15 concentration–response (also referred to as dose-response, DR) tokens combining assay context, log-concentration, and normalized response. The encoder is pretrained with masked-response reconstruction plus low-weight auxiliary heads for outcome and AC_50_ prediction. For probe evaluation, the backbone is frozen and linear heads for three-class outcome, binary active/inactive, and AC_50_ regression are trained on training-split embeddings and evaluated on held-out compounds. The same training configuration is then applied to the full dataset to produce the deployable Tox21mer foundation model.

The assay transformer is a four-layer, eight-head Transformer encoder with a hidden dimension of 768, comprising 43.5 million parameters in total. Each of the six categorical assay descriptors is first embedded into a 32-dimensional vector and then projected to the model hidden dimension; these projected embeddings are summed to form an assay-context vector. This context vector is added both to a learned CLS token and to each of the 15 concentration-response (also referred to as dose-response, DR) tokens. Each dose-response token also contains an MLP-based embedding of the log-transformed concentration and an MLP-based embedding of the normalized response value. Missing dose positions are represented with masking.

The sequence of 16 tokens ([CLS] + 15 dose tokens) was processed by a four-layer transformer encoder with eight attention heads per layer, hidden dimension 768, feed-forward dimension 3,072, and dropout 0.1 (**Table 2**).

**Table 2.**
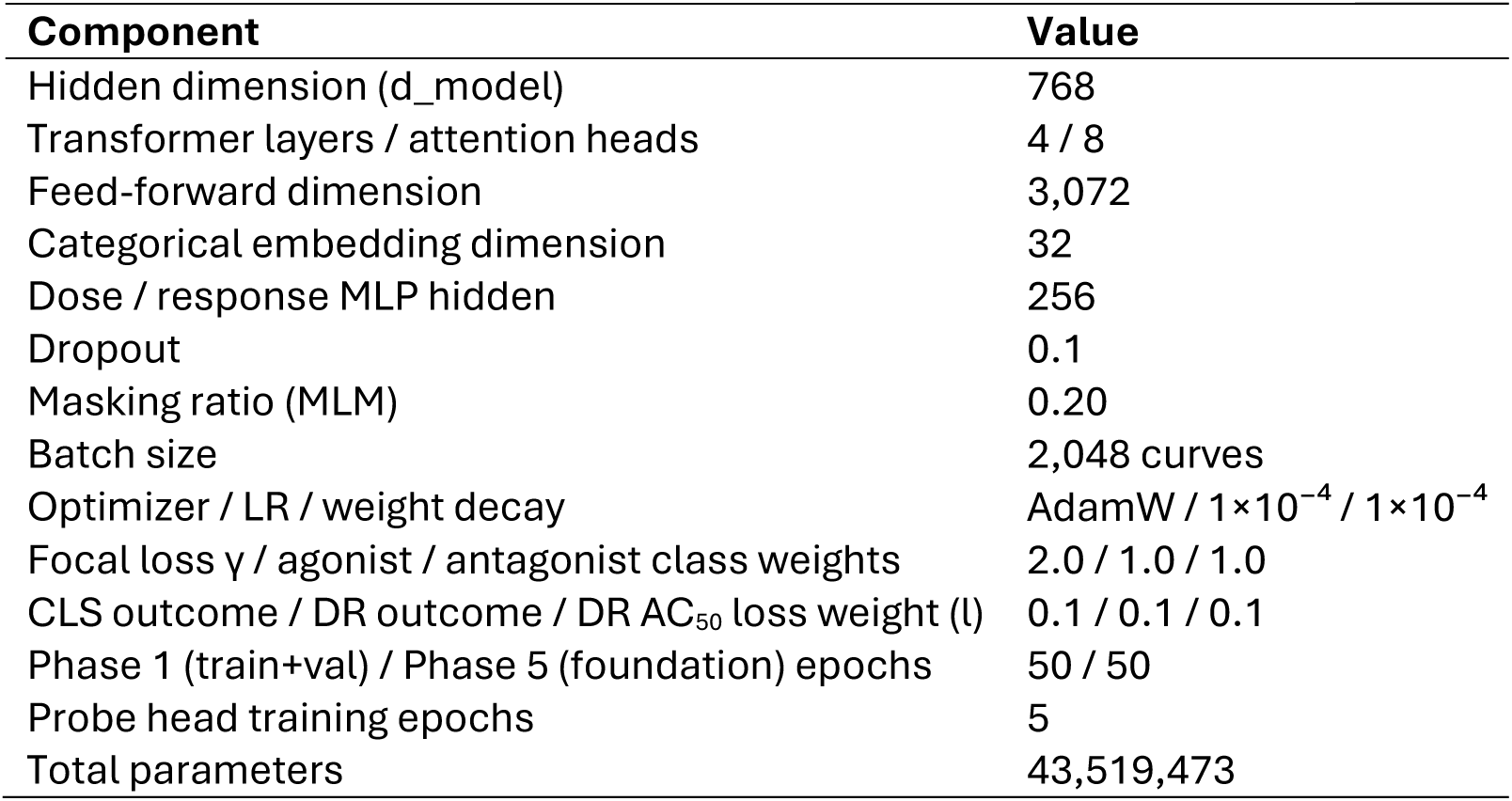
Tox21mer architecture and training hyperparameters.

### Embeddings

Each categorical descriptor was first mapped to a learned embedding of dimension (D=256) and then projected to the transformer hidden dimension (D=768). The projected embeddings from the six fields were summed to produce a single assay-context vector (ctx),

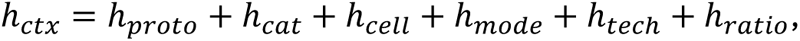

where proto encodes protocol name, cat encodes target category, cell refers to the cell line used in the assay, mode encodes assay mode, tech encodes the technology, and ratio indicates whether the curve corresponds to a ratio-based readout.

Each dose-response position was represented by a concentration-response token formed as *h_dr,i_* = *h_dose,i_* + *h_resp,i_*, where *h_dose,i_* is the embedding of the log-transformed concentration at position *i*, and *h_resp,i_* is the embedding of the corresponding normalized response value.

This ctx was used in two ways. First, it was added to a learned CLS (class) base token to form the sequence-level representation input. Second, it was added to each of the 15 dose-response tokens so that every token was explicitly conditioned on assay identity and experimental context.

The full input sequence was constructed by concatenating the context-conditioned CLS token with the 15 dose-response tokens, yielding a sequence length of 16. This sequence was passed through a Transformer encoder comprising 4 layers and 8 attention heads with hidden dimension 768, dropout, and a feedforward dimension of (4D). The model produced a contextualized output for every token, including a sequence-level representation from the CLS position and contextualized representations for all dose positions.

### Prediction heads

During backbone pretraining, task-specific heads were attached to the transformer outputs and optimized jointly. First, for masked-response reconstruction, a linear head was applied to each dose-response token to predict the masked response values. Second, for branch-specific outcome classification from the [CLS] embedding, linear heads were applied to the final [CLS] representation to predict outcomes for the mechanism and non-mechanism branches. Third, for auxiliary branch-specific outcome classification, additional linear heads were applied to the mean-pooled representation of the 15 dose-response tokens to predict the same mechanism and non-mechanism outcomes, with each auxiliary loss weighted by 0.1. Fourth, AC_50_ regression: a linear layer was applied to the same mean-pooled dose-response representation to predict log_10_(AC_50_), also weighted by 0.1. Full model hyperparameters are provided in Table 2.

For downstream evaluation after Phase 1 backbone pretraining, we attached two linear branch-probe heads: a three-class mechanistic head (agonist, antagonist, inactive) and a two-class non-mechanistic head (active, inactive). These probe heads were trained separately from the pretraining heads and are described further in Supplementary materials.

### Model Training

The primary pretraining objective was masked response reconstruction. For each assay curve, a subset of dose-response positions was randomly selected and treated as unknown. In our implementation, 3 of the 15 dose positions were masked. The input sequence to the model therefore consisted of a CLS token and 15 dose slots, in which the dose embeddings remained visible for all positions, while the response embeddings at masked positions were replaced by a learned mask embedding.

After passing the sequence through the stacked transformer encode, the final-layer hidden stages for curve *i*, denoted as 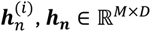, where *M*=16 corresponding to the CLS plus 15 concentration(dose)-response tokens, and *D* is the hidden dimension. A shared MLP was applied to the token representation to predict the log_10_-transformed response values, 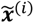, at each dose position *j*. The reconstruction loss was computed over the masked positions.

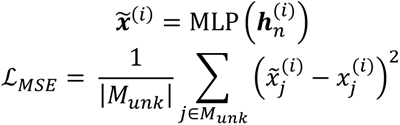

The total training loss also included a small contribution from the outcome-classification objective, weighted by λ = 0.1:

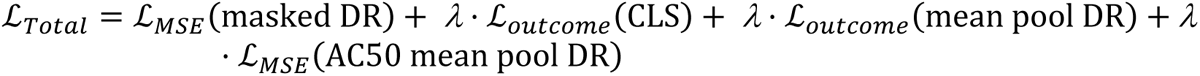

Where MSE is mean-squared error on masked positions, and *L_outcome_* is the focal loss *FL*(*p_t_*) = −*α_t_*(1 − *p_t_*)*^γ^ log*(*p_t_*) with *γ* = 2.0, where *p_t_* is the predicted probability assigned to the true outcome class and the class weights were set to and *a_t_* = 1.0 for agonist and antagonist. We adopted γ = 2.0 from Lin et al. (2017) as a starting hyperparameter rather than as the result of a tuned sweep. The non-mechanism head used plain cross-entropy.

A linear prediction head was applied to each dose-position output to predict the corresponding normalized response value. An additional scalar AC_50_ head was also instantiated for auxiliary prediction. Thus, the transformer jointly learned token-level representations of concentration–response structure and a sequence-level representation of assay-level activity.

The model was parameterized by the categorical embedding size, transformer hidden size, multilayer perceptrons (MLP) hidden sizes for dose and response encoding, number of encoder layers, number of attention heads, and dropout rate. The dose encoder mapped a scalar concentration value to a (d)-dimensional embedding through a linear layer, ReLU (Rectified Linear Unit) activation, and output projection. The response encoder used the same architecture to map a scalar response value to a (d)-dimensional embedding.

### Evaluation metrics

We report macro-F1 for the mechanistic head over the three classes (agonist, antagonist, inactive), binary F1 for the non-mechanistic head, and a combined score defined as 0.5 ×(mech_F1 + nonmech_F1). Full per-class precision, recall, and F1 values are provided in the Supplementary Information. For AC_50_ prediction, we report RMSE (root mean square error), MAE (mean absolute error), correlation R^2^, and Pearson correlation, both overall and on a per-protocol basis. For masked-response reconstruction, the primary metric is validation masked-position MSE.

### Computing

All training runs were performed on a single NVIDIA A100-SXM4 GPU (40 GB or 80 GB memory), without distributed training. Data loading used 8–16 worker processes depending on the run. A single Phase 1 pretraining run (50 epochs, 1,723,903 training rows, batch size 2,048) required approximately 10 hours. The probe-training phases (Phases 2–4) completed in approximately 15 minutes on a single GPU, whereas each MLP baseline finished in less than 2 hours. Phase 5 foundation-model retraining required an additional 10 hours.

### Ablation Study

We conducted extensive ablation studies to determine which aspects of the model and input representation drove downstream performance. These experiments included backbone, probing, and testing settings, each with the corresponding pretrained backbone kept fixed.

Block 1: probe-training and test-input perturbations with a clean backbone. we trained probe heads on either unaltered curves or curves in which the normalized response values were randomly permuted within each curve. Each probe was then evaluated on both clean and within-curve-shuffled test inputs, yielding a (2 × 2) design that separated the effects of the probe-training distribution from those of the test-time input structure.

Block 2: probe-training and test-input perturbations with a shuffled-data backbone. We repeated the same (2 × 2) probe analysis using a backbone that had itself been pretrained on curves with within-curve response shuffling applied dynamically during training and validation. All other pretraining and probe-training settings were kept unchanged.

Block 3: MLP baselines without attention. To measure how much predictive signal could be captured without a transformer, we trained two-layer MLP baselines on the same data splits using the same branch-specific setup, losses, label masking, and evaluation metrics as the transformer probes. We tested six input variants: summary statistics alone or with assay context, the raw 15-dimensional normalized response vector alone or with context, and permuted versions of the raw vector with or without context. These comparisons were intended to clarify what the supervised probe metric captures, rather than to evaluate the broader suitability of transformer representations for downstream tasks such as distillation, retrieval, or compound-level prediction. Additional methodological details are provided in the Supplementary Materials.

## Results

Training converged quickly and remained stable over 50 epochs (**Figure 2**). Masked-response reconstruction loss dropped sharply early on, with train masked MSE falling from 1.65 at epoch 1 to 0.54 at epoch 2 and gradually to 0.43 by epoch 50. Validation masked MSE followed a similar pattern, settling around 0.42–0.44 with little sign of overfitting. Validation outcome accuracy was already high after epoch 1 (0.946), exceeded 0.96 by epoch 3, and remained near 0.97 thereafter. Overall, the model learned reconstruction and label-relevant features within the first few epochs, with later training yielding only modest gains.

**Figure 2.**
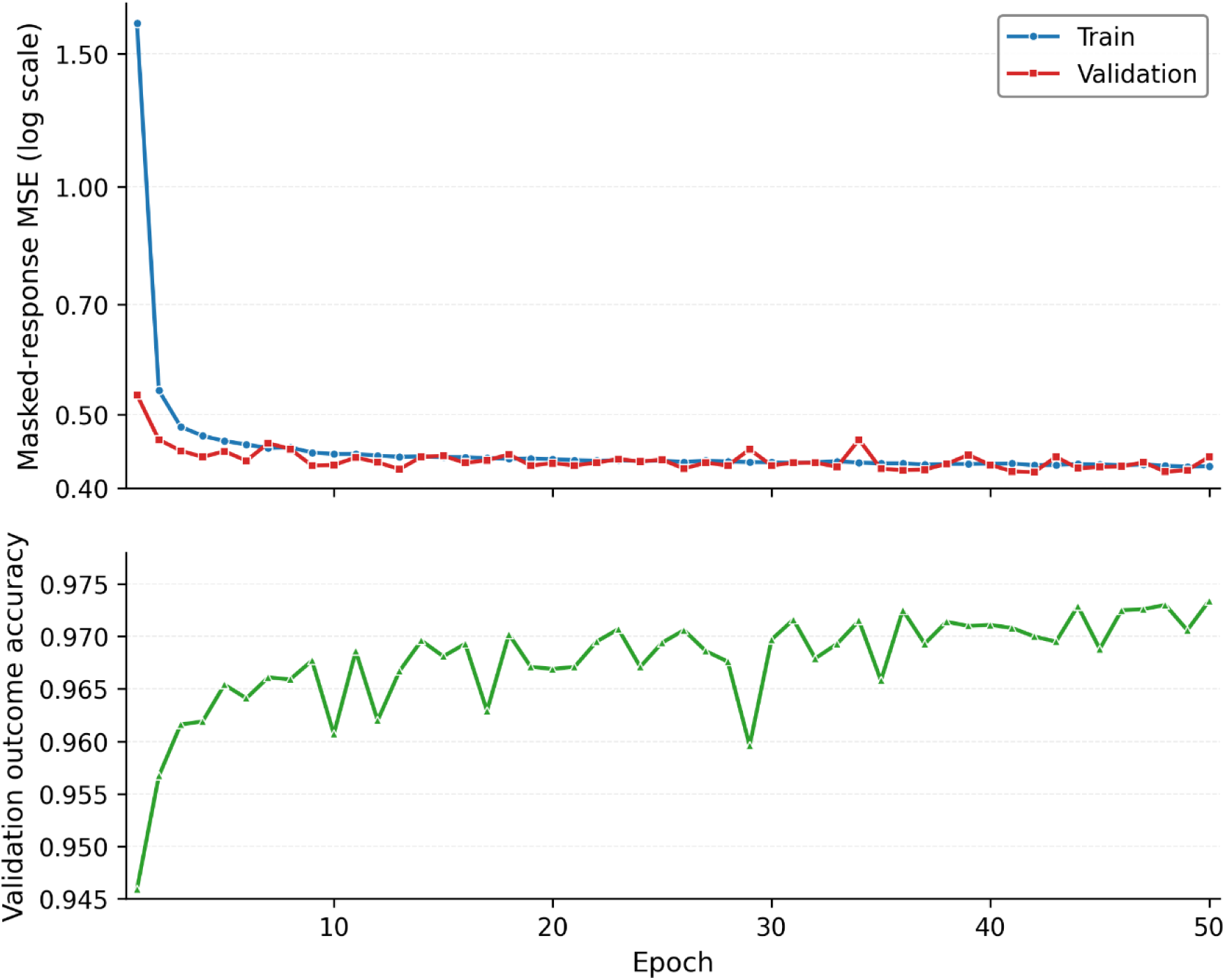
Phase 1 pretraining convergence. Per-epoch training and validation masked-response MSE (top, log scale) and validation outcome-classification accuracy (bottom) over 50 epochs of Phase 1 pretraining.

### Frozen embeddings support accurate outcome and AC_50_ prediction

All performance evaluations reported here were conducted on curves in the cc2 strict set; corresponding results for cc2 loose are provided in the Supplementary Materials (Table S2, S3 and Figure S2, S3). On held-out test compounds, the mechanistic branch comprised 9,302 agonist, 7,180 antagonist, and 147,746 inactive curves, while the non-mechanistic branch comprised 19,100 active and 65,651 inactive curves. The baseline model achieved a mechanistic macro-F1 of 0.985, with class-specific F1 scores of 0.978 for agonists, 0.979 for antagonists, and 0.998 for inactives. On the non-mechanistic branch, it achieved a binary F1 of 0.994, with class-specific F1 scores of 0.978 for actives and 0.994 for inactives (**Table 3a**). The confusion matrices are shown in **Figure 3**.

**Figure 3.**
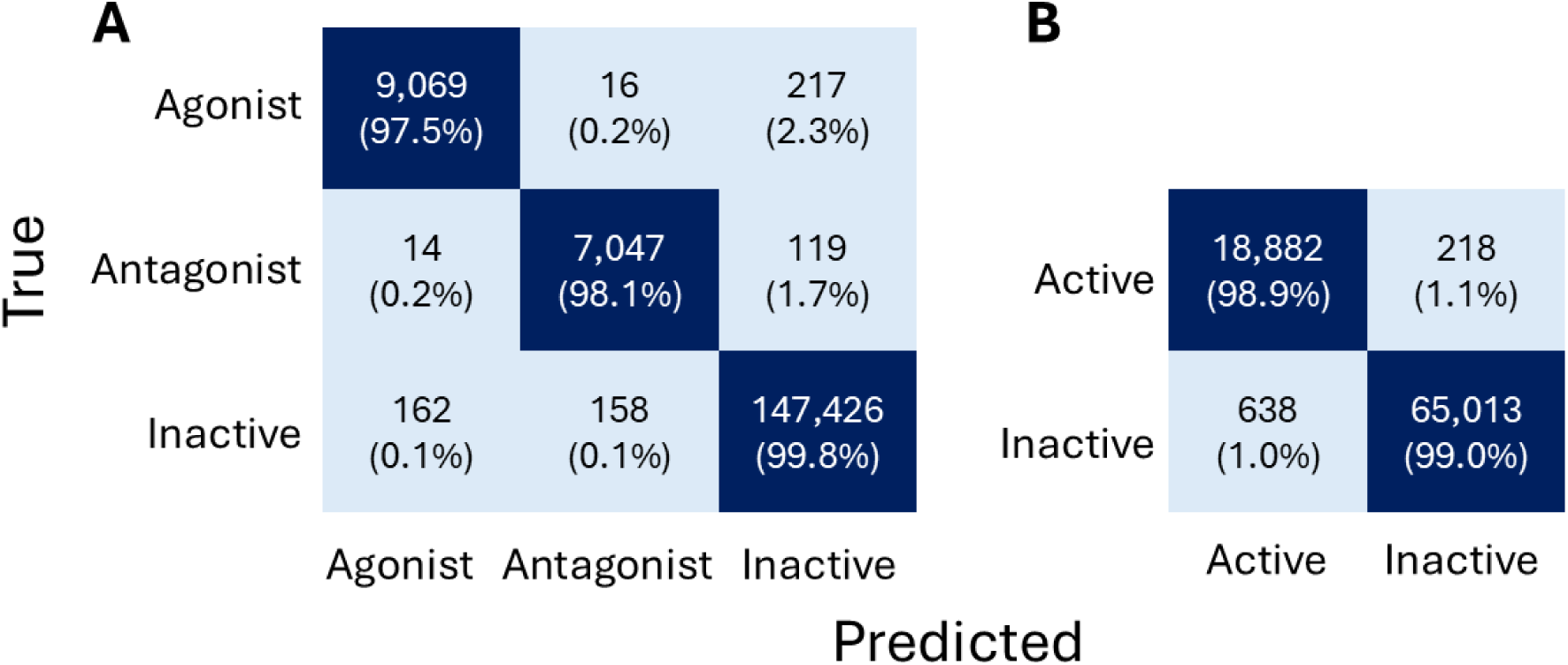
Confusion matrices for the mechanism and non-mechanism branch probes on held-out test compounds. Results are from a backbone trained with light auxiliary supervision and `cc2=strict`. (A) Three-class mechanism branch (n = 164,228); (B) Binary non-mechanism branch (n = 84,751).

**Table 3a.**
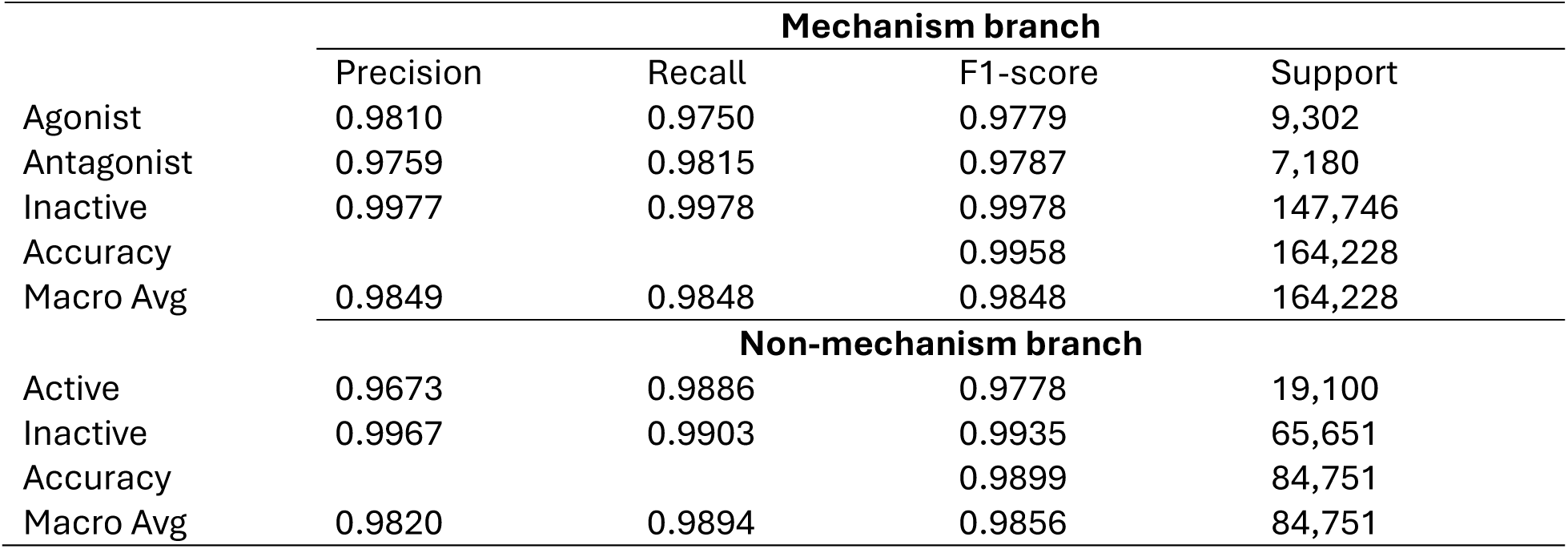
Test data mechanism and non-mechanism branch classification reports.

|  | Mechanism branch |  |  |  |
| --- | --- | --- | --- | --- |
|  | Precision | Recall | F1-score | Support |
| Agonist | 0.9810 | 0.9750 | 0.9779 | 9,302 |
| Antagonist | 0.9759 | 0.9815 | 0.9787 | 7,180 |
| Inactive | 0.9977 | 0.9978 | 0.9978 | 147,746 |
| Accuracy |  |  | 0.9958 | 164,228 |
| Macro Avg | 0.9849 | 0.9848 | 0.9848 | 164,228 |
|  | Non-mechanism branch |  |  |  |
|  | Precision | Recall | F1-score | Support |
| Active | 0.9673 | 0.9886 | 0.9778 | 19,100 |
| Inactive | 0.9967 | 0.9903 | 0.9935 | 65,651 |
| Accuracy |  |  | 0.9899 | 84,751 |
| Macro Avg | 0.9820 | 0.9894 | 0.9856 | 84,751 |

**Table 3b.** Per-class PR-AUC and ROC-AUC on the held-out test set.

|  | Branch Class | PR-AUC | ROC-AUC | Support |
| --- | --- | --- | --- | --- |
| mechanism | Agonist | 0.9976 | 0.9998 | 9,302 |
|  | Antagonist | 0.9973 | 0.9998 | 7,180 |
|  | Inactive | 1.0000 | 0.9997 | 147,746 |
| Non-mechanism | Active | 0.9981 | 0.9993 | 19,100 |

Precision–recall area under the curve (PR-AUC) was uniformly high across both branches, including the minority classes most relevant for downstream prioritization: 0.998 for agonists, 0.997 for antagonists, 1.0 for mechanistic inactives, and 0.998 for non-mechanistic actives. Confusion matrices for both branches (**Table 3b**) further show that the few remaining errors are concentrated in false negatives among the minority active classes, with little systematic confusion between agonist and antagonist.

For AC_50_ prediction across the same 102 protocols and 52,417 test-set evaluations, the unweighted regression probe showed strong agreement between predicted and observed log_10_(AC_50_) values, with a sample-weighted median per-protocol R^2^ of 0.88 and median Pearson correlation of 0.95, respectively. However, the scatter plot revealed systematic underestimation of potency for the most potent (lower AC_50_) compounds (low tail MAE = 0.44, low tail residual mean = 0.404). This bias is consistent with the right-skewed label distribution, in which very low AC_50_ values are underrepresented during training.

To mitigate this effect, we replaced the standard MSE objective with a weighted MSE loss, which substantially improved performance for the underrepresented high-potency compounds (**Figure 4**) with low tail MAE of 0.257 (compared to 0.44) and residual mean of 0.01 (compared to 0.404). Across all test curves, both the Pearson correlation between observed and predicted log_10_(AC_50_) were 0.94 and a R^2^ of 0.87. The strongest AC_50_ performances were observed for tox21-pr-bla-agonist-p1 and tox21-er-luc-bg1-4e2-agonist-p4 (**Figure 5**). In contrast, the weakest performances were observed for tox21-fxr-bla-agonist-p2 and tox21-sbe-bla-agonist-p1 (**Figure 5**). The AC_50_ performance ranked by R^2^ is provided in Supplementary **Table S4**. Relatively weak performance was also observed for tox21-kiss1r-wt assay.

**Figure 4.**
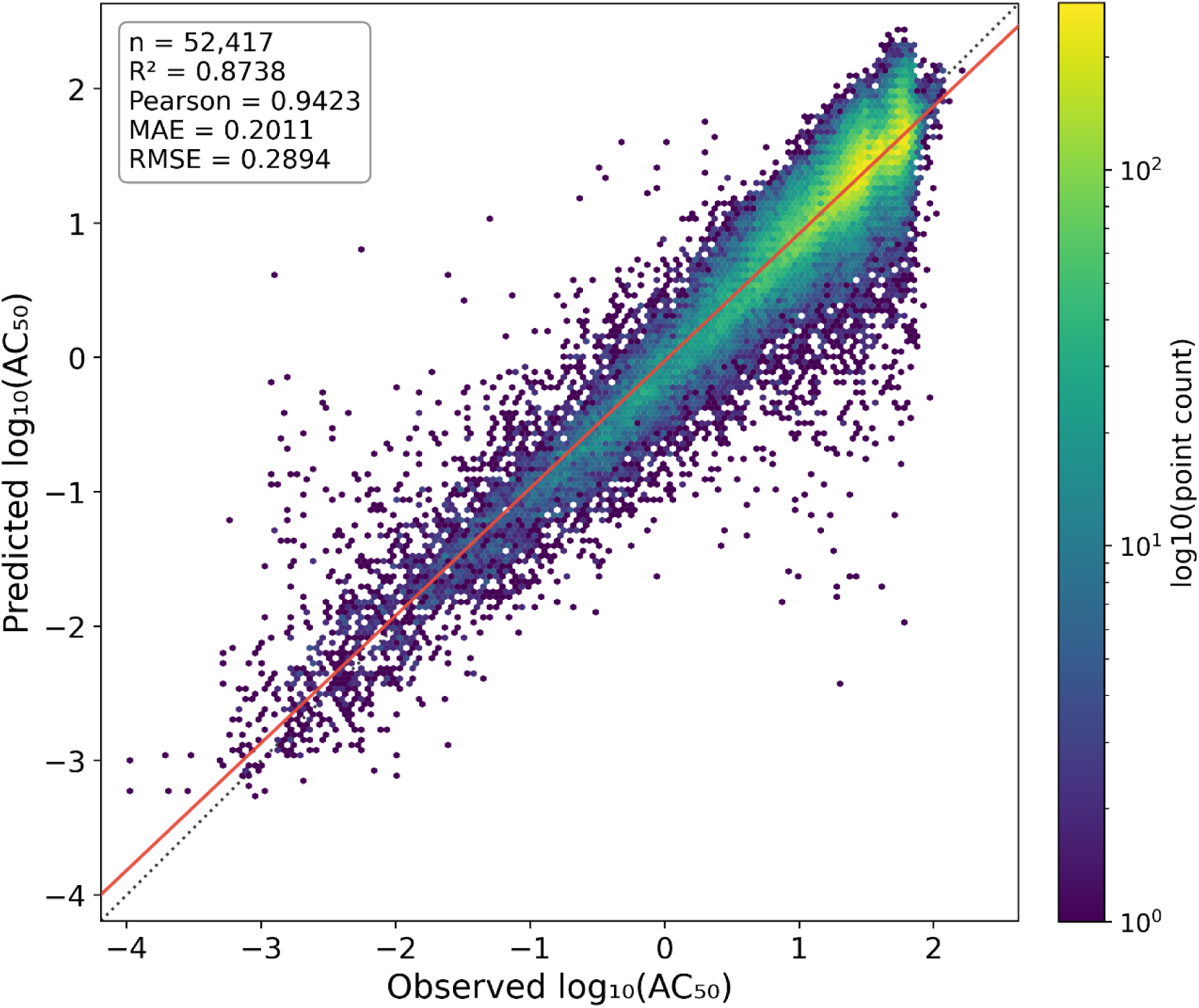
Predicted log_10_(AC_50_) vs observed log_10_(AC_50_) values for all cc2=strict test curves. The dotted line indicates the identity line, and the solid red line shows ordinary-least-squares fitted line.

**Figure 5.**
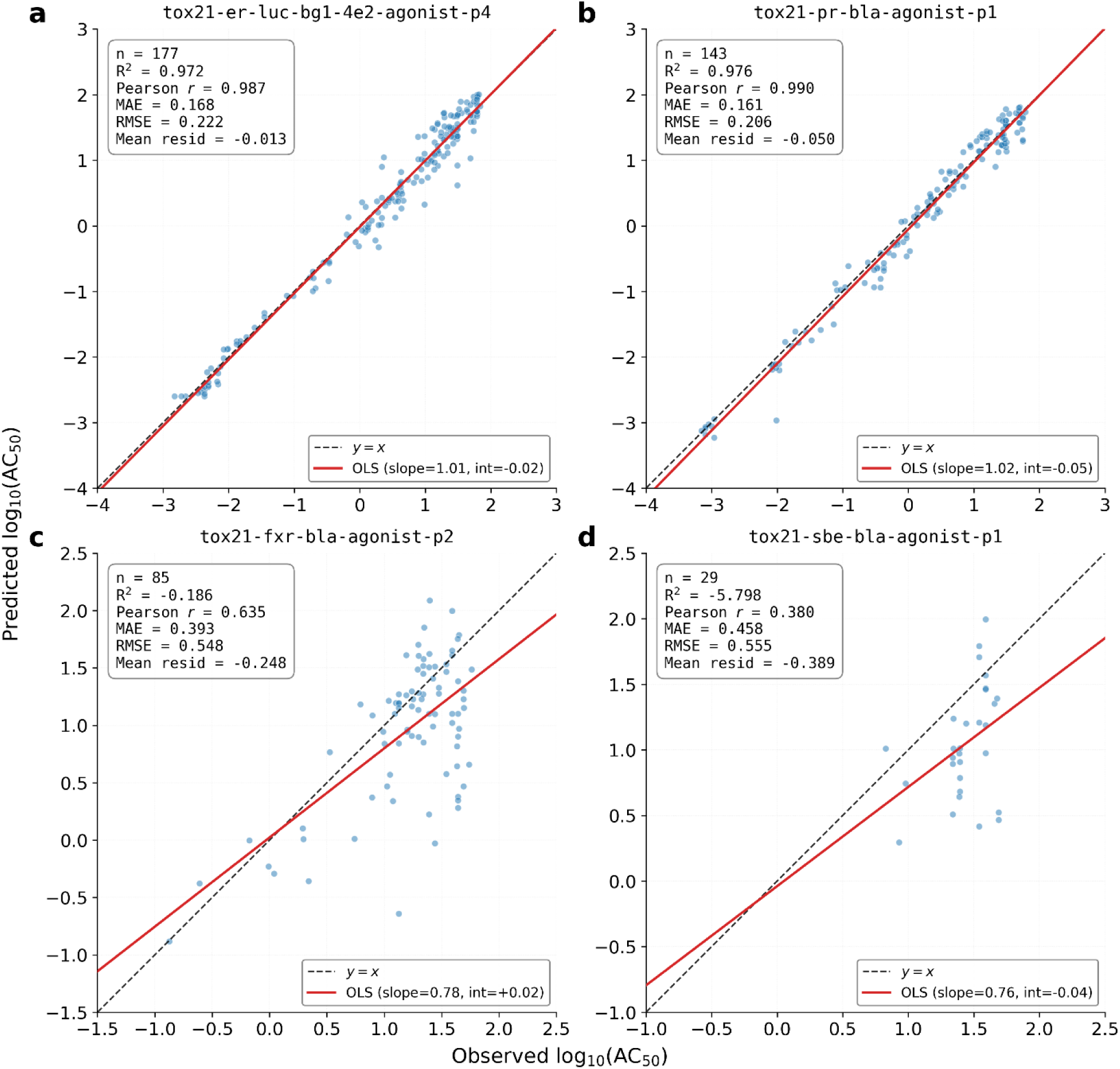
Predicted versus observed log_10_(AC_50_) for two best and two worst protocols. The ordinary-least-squares fit (red) overlaid on the y = x reference (dashed). (a) tox21-er-luc-bg1-4e2-agonist-p4. (b) tox21-pr-bla-agonist-p1. (c) t tox21-fxr-bla-agonist-p2. (d) tox21-sbe-bla-agonist-p1.

AC_50_ prediction performance varied across the 102 protocols. Antagonist-labeled assays showed a trend toward better AC_50_ prediction than agonist-labeled assays, with mean and median R^2^ values of 0.834 and 0.857, respectively, compared with 0.596 and 0.796 for agonist-labeled assays (*P* = 0.09, two-sided Mann–Whitney *U* test). For this comparison, we excluded the three assays with mode of “agonist + antagonist”. No differences were found between mechanism and non-mechanism branch (median R^2^ = 0.825 vs 0.821, *P*=0.6, two-sided *U* test).

### The pretrained embeddings capture curve-class structure

We examined whether the model’s pooled sequence representation (CLS embedding) captured information related to assay curve class by grouping each compound–assay instance according to its cc2 label into three predefined categories: high-quality positive curves, high-quality negative curves, and flat inactive curves (cc2 = 4). For each retained row, we extracted the corresponding CLS vector and projected the embeddings into two dimensions using t-SNE, with points colored by cc2 category. The resulting visualizations showed partial separation and coherent local grouping of these categories, consistent with the interpretation that the learned embedding retains information associated with curve-class structure (**Figure 6**).

**Figure 6.**
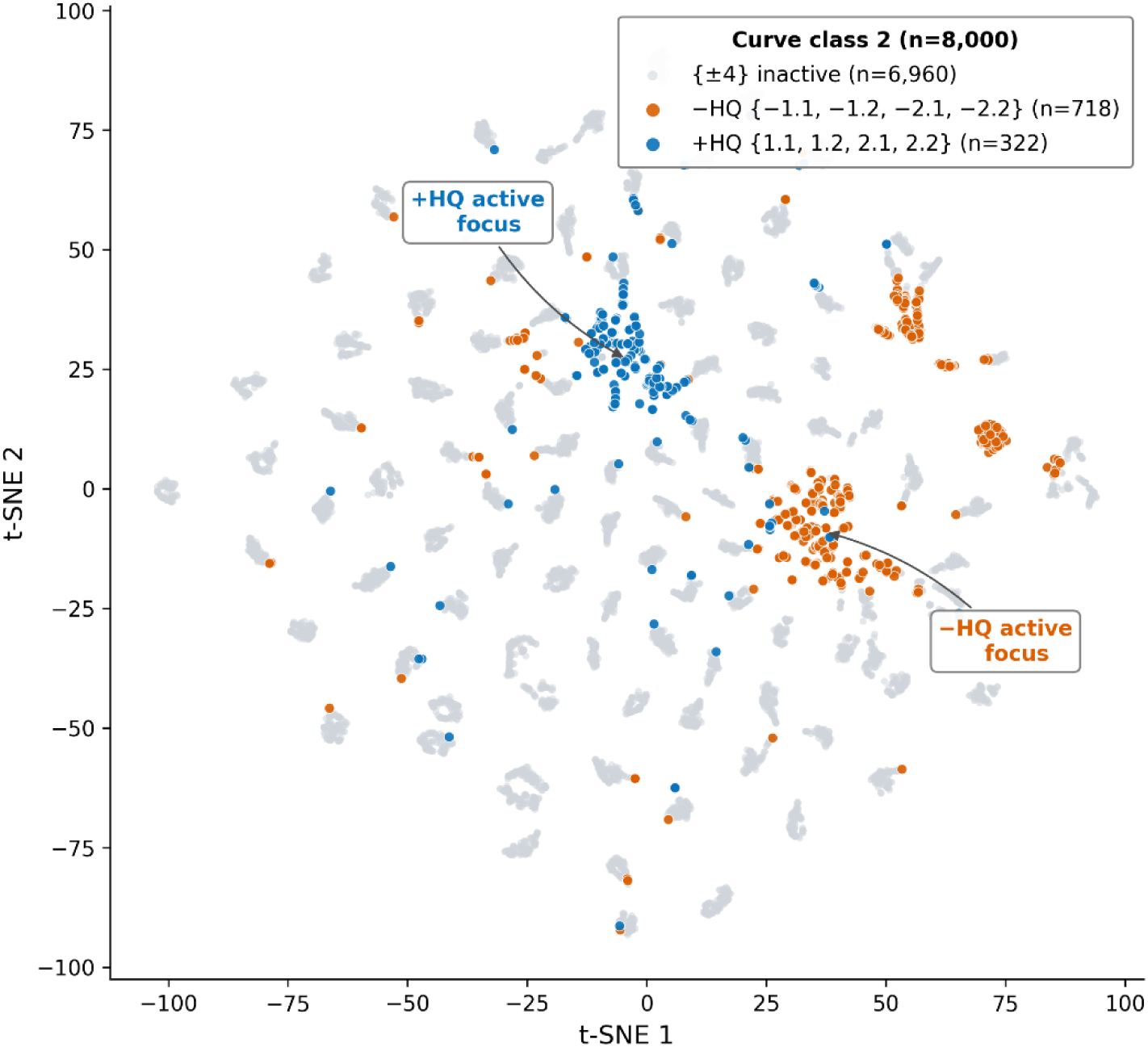
CLS embeddings cluster by Curve_class2 quality category. CLS embeddings cluster by cc2 category. Only a stratified subsample of 8,000 of the 529,701 unique embeddings is shown.

### Self-supervised pretraining alone yields a strong representation

Because Tox21mer pretraining includes lightly weighted auxiliary outcome and AC_50_ losses (λ = 0.1) that overlap with the downstream probe labels, probe F1 could partly reflect retained pretraining supervision rather than representation quality. To test this, we retrained Tox21mer from the same initialization with all auxiliary losses removed, so Phase 1 and foundation training used only masked-response reconstruction under the same data, splits, and hyperparameters; branch probes were then trained as before on the frozen self-supervised backbone.

Without auxiliary supervision, Tox21mer still performed strongly, reaching mechanism macro-F1 0.957 (agonist 0.925, antagonist 0.952, inactive 0.994) and non-mechanism binary-F1 0.986 (active 0.947, inactive 0.986). PR-AUC remained high for all classes, exceeding 0.977 throughout. Relative to the auxiliary-supervised model (**Table S5**), auxiliary losses improved F1 by 1–6 points, with the largest gains on minority active classes (agonist +5.3, active +3.1), suggesting modestly sharper biologically relevant decision boundaries beyond masked-response reconstruction alone. These results indicate that most probe performance arises from the masked-response objective rather than simple retention of auxiliary labels. We therefore interpret the probe metrics as reflecting generalization across held-out compounds rather than memorization of the auxiliary task family. We adopt the auxiliary-supervised configuration as the deployable Tox21mer model because its representation is better aligned with the intended downstream use cases, including external chemical prediction and distillation into chemistry-only student models for outcome and AC_50_ prediction. In this setting, low-weight auxiliary supervision provides useful prior information about activity and potency while preserving a backbone that remains informative even when trained with masked-response reconstruction alone.

### Predictive performance is driven primarily by curve-level response-value distributions

With the clean-trained backbone (**Table 4**, rows 1–4), the fully matched clean condition (row 1) achieved a mechanistic F1 of 0.9848. In contrast, when the same probe was evaluated on shuffled test curves (row 2), F1 score dropped sharply to 0.5275, largely due to a collapse in recall for the active classes (0.202 and 0.174, for agonist and antagonist). However, when the probe itself was trained on shuffled curves and evaluated on shuffled inputs (row 3), performance recovered substantially to 0.9273. Evaluation of that shuffled-trained probe on clean test curves (row 4) also remained high, with an F1 of 0.9.

**Table 4.** Results of Ablation Studies. Mechanism and non-mechanism F1 across eight probe conditions.

| Phase | Backbone | Probe DR | Test DR | Mech F1 | Non-mech F1 |
| --- | --- | --- | --- | --- | --- |
| 2a | clean | clean | clean | 0.9848 | 0.9935 |
| 2b |  | clean | shuffled | <u>0.5275</u> | 0.9011 |
| 2c |  | shuffled | shuffled | 0.9273 | 0.9801 |
| 2d |  | shuffled | clean | 0.9000 | 0.9726 |
| 2a | shuffled | clean | clean | 0.9494 | 0.9883 |
| 2b |  | clean | shuffled | <u>0.7823</u> | 0.9655 |
| 2c |  | shuffled | shuffled | 0.9097 | 0.9813 |
| 2d |  | shuffled | clean | 0.9103 | 0.9806 |

Repeating the same four conditions with a backbone pretrained on shuffled curves (**Table 4**, rows 5–8) reproduced the qualitative pattern observed for the clean backbone, with two informative shifts. First, the matched shuffled-backbone / clean-probe / clean-test condition (row 5) still achieved a mechanistic F1 of 0.9494 and a non-mechanistic F1 of 0.9883, only ~3 points below the clean-backbone baseline, demonstrating that pretraining on randomized within-curve orderings does not impair the backbone’s predictive utility on naturally ordered curves. Second, the clean-probe / shuffled-test mismatch (row 6) recovered substantially, from 0.5275 on the clean backbone to 0.7823 on the shuffled backbone. Taken together, these results show that probe-training distribution mismatch has a larger effect on downstream performance than whether the backbone was pretrained on ordered or shuffled response sequences. The remaining asymmetry between Phases 2b and 2d also decreases when the backbone is exposed to a wider range of input orderings during pretraining. A separate ablation indicated that assay context is only weakly informative for these probe tasks (Supplementary **Table S6**).

To verify that the curve-shape conclusion does not depend on the transformer architecture, we additionally trained two-layer MLPs (256 hidden units, GELU, dropout 0.2) on the same train/validation/test splits with the same branch-probe loss structure and evaluation metrics, replacing the transformer backbone with explicit feature representations of the response values. Six configurations were evaluated, varying whether the input was eight hand-crafted summary statistics or the raw 15-dimensional response vector, whether assay-context features were included, and whether the raw response vector was randomly permuted within each curve at every training step.

Both raw-input and randomly-permuted-input MLPs reached comparable F1 to the transformer when assay context was included, again with no advantage from preserving within-curve order (Supplementary **Table S7**). The MLP comparison is intended to test what the supervised probe metric is sensitive to — specifically that the metric does not require curve-shape information; it does not address the suitability of the transformer representation for downstream uses such as distillation, retrieval, or compound-level external screening, where embedding structure matters beyond probe accuracy.

### Foundation model

To produce a deployable model for downstream use, we retrained Tox21mer on the full quality-controlled dataset, comprising 2,489,975 curves from 6,727 compounds with unique SMILES ^31^ (6,743 unique CAS numbers; 9,693 unique Tox21 IDs) across 102 protocols. This full-data run used the same architecture, optimizer, loss configuration, batch size (2,048), and 50-epoch training schedule as the split-based Phase 1 pretraining. Because all original train, validation, and test rows were included at this stage, no held-out set remained for model selection; the final-epoch checkpoint was therefore saved as the Tox21mer foundation model, while unbiased estimates of generalization remain those reported from the original split-based evaluation.

To support application to new Tox21-format data, we saved the foundation-model weights together with the preprocessing artifacts required for inference, including categorical vocabularies, per-protocol response winsorization and normalization statistics, and log-dose standardization parameters. New curves processed through this same pipeline can be embedded directly into a 768-dimensional CLS representation, providing a fixed-length, context-aware encoding of the full dose–response profile. Per-dose token embeddings are also available when finer-grained analyses are needed.

### Potential utilities of Tox21mer

In practical terms, Tox21mer serves as a foundation model by providing pretrained, reusable embeddings for Tox21 concentration–response data. Rather than being optimized for a single assay in isolation, the model is trained across the full Tox21 panel and therefore learns a shared, assay-aware representation of response patterns in context. These embeddings may support a range of downstream applications, including compound similarity analysis, clustering of response profiles, retrieval of biologically related curves, and transfer learning to new endpoints or related screening datasets. The framework may also be useful for assay quality assessment and for the imputation or denoising of partially observed concentration–response measurements, where a pretrained representation of typical response behavior could be beneficial. More broadly, the embeddings provide a common representation for comparing, organizing, and interpreting large-scale qHTS data across protocols, compounds, and biological targets. With appropriate downstream adaptation, Tox21mer may also support prospective screening of external chemicals across Tox21 endpoints, with potential applications in drug safety assessment and computational hazard prioritization.

## Discussion

We developed Tox21mer, a transformer-based foundation model for Tox21 quantitative high-throughput screening data that jointly encodes a compound’s 15-point concentration–response curve and its assay-context metadata into a single 768-dimensional representation. The model was trained on approximately 2.5 million curves spanning 102 protocols and 6,727 unique compounds. In probe-based evaluation on curves from held-out compounds, the learned representation supported strong performance on both a three-class outcome task (agonist, antagonist, inactive) and a binary active/inactive task. It achieved a macro-F1 of 0.985 on the mechanistic task and a binary F1 of 0.995 on the non-mechanistic task. The model also provided competitive AC_50_ prediction with a sample-weighted median per-protocol R^2^ of 0.87 across the 83,143 test-set evaluations. The t-SNE projection of CLS embeddings additionally showed coherent local grouping by curve-class category, consistent with the interpretation that the learned representation retains curve-quality structure. These results establish Tox21mer as a reusable, assay-aware backbone for Tox21 data.

A light-weight auxiliary-supervision scheme was retained in the final Tox21mer training objective for a practical reason: the intended downstream use of the backbone is not generic representation learning in the abstract, but external chemical prediction and distillation into chemistry-only student models for outcome and AC_50_ prediction. In that setting, light-weighted supervised signals on assay outcome and potency provide useful task-aligned inductive bias, encouraging the backbone to organize its latent space around biologically relevant activity and potency axes while leaving masked-response reconstruction as the dominant training objective. The masked-only control shows that the representation is already highly informative without auxiliary labels, whereas the auxiliary-supervised variant yields consistent gains, concentrated on the minority active classes and potency-related structure most relevant for downstream prioritization. We therefore view the auxiliary terms not as the primary source of performance, but as a light regularizing prior that improves downstream utility while preserving the broad unsupervised structure learned from the full corpus of concentration–response curves.

The Tox21 data are heterogeneous, so we applied four distinct preprocessing steps. First, the protocols included both mechanism-based assays (agonist, antagonist) and non-mechanism assays (e.g., cell viability, cytotoxicity), which made it difficult to use a single outcome head. We therefore split the assays by assay mode into mechanism and non-mechanism branches. This enabled branch-specific prediction and avoided inappropriate predictions, such as assigning agonist or antagonist labels to cell-viability assays. Second, some assay datasets contained multiple modalities. For example, the DT40 dataset includes mitochondrial cytotoxicity outcomes for three different cell types. Similarly, two cell-viability assays included multiple time points, each with its own outcome, for the same protocol–chemical combination. Without separating these into individual datasets, the model would encounter multiple different outcomes for the same protocol–chemical pair, creating ambiguity. Third, mechanism labels (agonist and antagonist) were sometimes assigned to curves from cytotoxicity assays. To reduce confusion, we reassigned non-mechanism assay outcomes solely on the basis of curve class, using the labels active, inactive, or unknown. Fourth, we excluded curves labeled as “inconclusive”. However, we retained labels such as “inconclusive agonist.” We reasoned that it would not be meaningful to predict an inconclusive category when the experiment itself had failed to produce a robust response profile. These design choices made both model training and evaluation feasible.

For AC_50_ prediction, the unweighted regression probe showed strong agreement between predicted and observed log_10_(AC_50_) values. However, scatter plots revealed a systematic underestimation of potency for the most potent compounds (those with lower AC_50_ values), likely due to the severe imbalance caused by many inactive curves with high AC_50_ values. To address this issue, we replaced the standard MSE loss in the AC_50_ head with a weighted MSE loss. This approach improved prediction not only for the most potent curves but also in terms of overall R^2^. We also observed that AC_50_ prediction performance varied across protocols. In general, predictive ability was high in all instances, exhibiting slightly higher accuracy for mechanism-based curves than for non-mechanism curves, and antagonist curves appeared to be somewhat easier to predict than agonist curves.

An interesting finding from the ablation analysis is that Tox21mer’s CLS representation is largely insensitive to within-curve response ordering, provided that the probe is trained on a matching input distribution. Backbones pretrained on shuffled curves retained near-baseline performance when evaluated with probes trained on similarly shuffled inputs, whereas probes trained on clean curves generalized poorly to shuffled test curves. This asymmetric drop is best explained by probe train–test mismatch rather than by loss of predictive information in the backbone. A parallel comparison with a non-attention multilayer perceptron showed the same pattern, with no benefit from preserving within-curve order. Together, these results indicate that, under the present supervision setting, predictive performance is driven mainly by the distribution of response values conditioned on assay context rather than by detailed curve shape. This reflects the structure of the task rather than a failure of the model: most labeled curves are inactive, agonist and antagonist assays typically produce opposite-signed responses, and protocol identity is strongly associated with mode of action, making response magnitude and assay metadata highly discriminative.

Consistent with our preliminary experiments, teacher–student distillation provided little additional benefit for downstream outcome prediction when the student was already trained with direct supervision. This suggests that chemical structure, protocol embeddings, and supervised labels already capture most of the information needed for this task, and/or that a strong teacher may be aligned to a different objective than simple outcome classification. We therefore speculate that such a teacher may be more valuable for curve-level tasks, particularly the prediction of concentration-response behavior and other assay properties that depend on response-shape information.

Our study has limitations. First, evaluation was based on random SMILES-level splits. Although this prevents exact identity leakage between training and test, structurally related analogues of training compounds may still appear in the test set, potentially inflating absolute performance relative to scaffold-based or temporal-split benchmarks. Generalization to chemically novel classes outside the Tox21 library remains to be evaluated directly using external screening datasets. Incorporating additional chemical descriptors, such as ToxPrint ^32^ and Morgan fingerprints ^33,34^, together with similarity assessment against the existing Tox21 chemical reservoir, may further improve external screening performance. Second, our ablation strategy removes information by permutation but does not introduce tasks that explicitly require concentration–response shape. More direct tests of shape-aware representations would include benchmarks based on Hill-slope recovery, inflection-point identification, or transfer across assay protocols with different concentration grids.

In conclusion, Tox21mer provides a practical foundation model for *in vitro* concentration-response data across the full Tox21 assay panel. By learning protocol-aware embeddings from large-scale qHTS response profiles, the model captures shared patterns of assay context and response behavior in a reusable latent space, rather than being optimized for any single assay in isolation. Tox21mer may support a range of downstream applications, including compound similarity analysis, clustering and retrieval of biologically related response profiles, transfer learning to new endpoints or related screening datasets, assay quality assessment, and the imputation or denoising of partially observed concentration-response measurements. More broadly, Tox21mer offers a common representation for comparing, organizing, and interpreting qHTS data across protocols, compounds, and biological targets, and, with appropriate downstream adaptation, may support prospective screening of external chemicals across Tox21 endpoints for drug safety assessment and computational hazard prioritization.

## Code and data Availability

Raw Tox21 screening data are available from the NIH Toxicology in the 21st Century program (https://tripod.nih.gov/tox21/). The QC-filtered corpus, splits, vocabularies, normalization statistics, and all trained checkpoints used in this study are available at https://github.com/yuanyuanli66/tox21-assay-backbone.

## Author contributions

LL conceived, designed, and conducted the study, performed all analyses, and wrote the initial draft of the manuscript. JH provided critical input on the study design and the initial draft. KS, AM-R, J-HH, SA, and DR contributed to study design and manuscript revision. YL assisted with the study and led code deposition.

## Declaration

Artificial intelligence tools were used to assist in code development and manuscript preparation.

## Acknowledgements

We are grateful to the NIEHS Office of Scientific Computing, especially Dr. Frank Day, for computing support. This research was supported in part by the Intramural Research Program of the National Institutes of Health (NIH), National Institute of Environmental Health Sciences (ZIA ES101765). The contributions of the NIH authors were made as part of their official duties as NIH federal employees, are in compliance with agency policy requirements, and are considered Works of the United States Government. However, the findings and conclusions presented in this paper are those of the authors and do not necessarily reflect the views of the NIH or the U.S. Department of Health and Human Services (HHS).

## Competing interests

The authors declare no competing interests.

## Notes

https://github.com/yuanyuanli66/tox21-assay-backbone

### Competing Interest Statement

The authors have declared no competing interest.

