## supplement for "Tox21mer, A transformer foundation model for Tox21 high-throughput concentration–response curves data"

**Table S1**. Tox21mer metadata.

| Protocol Name | Assay Target | Target Category | Cell Line | Cell Type | Mode | Technology |
|---|---|---|---|---|---|---|
| tox21-ache-p3 | AChE (colorimetric) | Neurotoxicity | SH-SY5Y | Neuroblast | Inhibition | Colorimetric |
| tox21-ache-p4 | AChE (fluorescent) | Neurotoxicity | SH-SY5Y | Neuroblast | Inhibition | Fluorescence |
| tox21-ache-p5 | AChE | Neurotoxicity | NA | Biochemical | Inhibition | Colorimetric |
| tox21-adrb2-agonist-p1 | ADRB2 agonist | GPCR | CHO | Hamster | Agonist | Fluorescence |
| tox21-adrb2-antagonist-p1 | ADRB2 antagonist | GPCR | CHO | Hamster | Antagonist | Fluorescence |
| tox21-ahr-p1 | AhR | NR | HepG2 | Liver | Agonist | Luciferase |
| tox21-ap1-agonist-p1 | AP-1 agonist | SR | ME-180 | Cervical Cancer | Agonist | Beta-lactamase |
| tox21-ar-bla-agonist-p1 | AR-BLA agonist | NR | HEK293 | Kidney | Agonist | Beta-lactamase |
| tox21-ar-bla-antagonist-p1 | AR-BLA antagonist | NR | HEK293 | Kidney | Antagonist | Beta-lactamase |
| tox21-are-bla-p1 | ARE | SR | HepG2 | Liver | Agonist | Beta-lactamase |
| tox21-ar-mda-kb2-luc-agonist-p1 | AR-MDA agonist | NR | MDA-MB-453 | Breast Cancer | Agonist | Luciferase |
| tox21-ar-mda-kb2-luc-agonist-p3 | AR-MDA agonist (with antagonist) | NR | MDA-MB-453 | Breast Cancer | Agonist | Luciferase |
| tox21-ar-mda-kb2-luc-antagonist-p1 | AR-MDA antagonist | NR | MDA-MB-453 | Breast Cancer | Antagonist | Luciferase |
| tox21-ar-mda-kb2-luc-antagonist-p2 | AR-MDA antagonist (lower agonist) | NR | MDA-MB-453 | Breast Cancer | Antagonist | Luciferase |
| tox21-aromatase-p1 | Aromatase | SR | MCF-7 | Breast Cancer | Inhibition | Luciferase |
| tox21-car-agonist-p1 | CAR agonist | NR | HepG2 | Liver | Agonist | Luminescence |
| tox21-car-antagonist-p1 | CAR antagonist | NR | HepG2 | Liver | Antagonist | Luminescence |
| tox21-casp3-cho-p1 | Caspase-3/7 | Cytotoxicity | CHO | Hamster | Activation | Luminescence |
| tox21-casp3-hepg2-p1 | Caspase-3/7 | Cytotoxicity | HepG2 | Liver | Activation | Luminescence |
| tox21-chrm1-p1 | CHRM1 agonist and antagonist | GPCR | CHO | Hamster | Agonist and Antagonist | Fluorescence |
| tox21-cre-agonist-p1 | CRE agonist | SR | HEK293 | Kidney | Agonist | Beta-lactamase |
| tox21-cre-antagonist-p1 | CRE antagonist | SR | HEK293 | Kidney | Antagonist | Beta-lactamase |
| tox21-drd2-agonist-p1 | DRD2 agonist | GPCR | HEK293 | Kidney | Agonist | Fluorescence |
| tox21-drd2-antagonist-p1 | DRD2 antagonist | GPCR | HEK293 | Kidney | Antagonist | Fluorescence |
| tox21-dt40-p1-100 | Cell viability | Gene Tox | DT40 | Chicken | Cytotoxicity | Luminescence |
| tox21-dt40-p1-653 | Cell viability | Gene Tox | DT40 | Chicken | Cytotoxicity | Luminescence |
| tox21-dt40-p1-657 | Cell viability | Gene Tox | DT40 | Chicken | Cytotoxicity | Luminescence |
| tox21-elg1-luc-agonist-p1 | ATAD5 | Gene Tox | HEK293 | Kidney | Agonist | Luciferase |
| tox21-erb-bla-antagonist-p1 | ER-beta antagonist | NR | HEK293 | Kidney | Antagonist | Beta-lactamase |
| tox21-erb-bla-p1 | ER-beta agonist | NR | HEK293 | Kidney | Agonist | Beta-lactamase |
| tox21-er-bla-agonist-p2 | ER-BLA agonist | NR | HEK293 | Kidney | Agonist | Beta-lactamase |
| tox21-er-bla-antagonist-p1 | ER-BLA antagonist | NR | HEK293 | Kidney | Antagonist | Beta-lactamase |
| tox21-er-luc-bg1-4e2-agonist-p2 | ER-BG1 agonist | NR | BG1 | Ovarian | Agonist | Luciferase |

| | | | | | | |
|---|---|---|---|---|---|---|
| tox21-er-luc-bg1-4e2-agonist-p4 | ER-BG1 agonist (with antagonist) | NR | BG1 | Ovarian | Agonist | Luciferase |
| tox21-er-luc-bg1-4e2-antagonist-p1 | ER-BG1 antagonist | NR | BG1 | Ovarian | Antagonist | Luciferase |
| tox21-er-luc-bg1-4e2-antagonist-p2 | ER-BG1 antagonist (lower agonist) | NR | BG1 | Ovarian | Antagonist | Luciferase |
| tox21-err-p1 | ERR | NR | HEK293 | Kidney | Agonist | Fluorescence |
| tox21-esre-bla-p1 | ER stress | SR | HeLa | Cervical Cancer | Agonist | Beta-lactamase |
| tox21-fxr-bla-agonist-p2 | FXR-BLA agonist | NR | HEK293 | Kidney | Agonist | Beta-lactamase |
| tox21-fxr-bla-antagonist-p1 | FXR-BLA antagonist | NR | HEK293 | Kidney | Antagonist | Beta-lactamase |
| tox21-gh3-tre-agonist-p1 | TR-beta agonist | NR | GH3 | Rat pituitary | Agonist | Luciferase |
| tox21-gh3-tre-antagonist-p1 | TR-beta antagonist | NR | GH3 | Rat pituitary | Antagonist | Luciferase |
| tox21-gnrhr-hek293-p2 | GnRHR agonist | GPCR | HEK293 | Kidney | Agonist | Fluorescence |
| tox21-gr-hela-bla-agonist-p1 | GR-BLA agonist | NR | HeLa | Cervical Cancer | Agonist | Beta-lactamase |
| tox21-gr-hela-bla-antagonist-p1 | GR-BLA antagonist | NR | HeLa | Cervical Cancer | Antagonist | Beta-lactamase |
| tox21-h2ax-cho-p2 | H2AX | Gene Tox | CHO | Hamster | Activation | Fluorescence |
| tox21-hdac-p1 | HDAC | Gene Tox | HCT-116 | Colon Cancer | Inhibition | Luminescence |
| tox21-herg-u2os-p1 | hERG | Cardiotoxicity | U2OS | Osteosarcoma | Inhibition | Fluorescence |
| tox21-hre-bla-agonist-p1 | HRE-BLA agonist | SR | ME-180 | Cervical Cancer | Agonist | Beta-lactamase |
| tox21-hse-bla-p1 | HSE-BLA | SR | HeLa | Cervical Cancer | Agonist | Beta-lactamase |
| tox21-htr2a-p1 | HTR2A agonist and antagonist | GPCR | CHO | Hamster | Agonist and Antagonist | Fluorescence |
| tox21-kiss1r-hek293-p2 | KISS1R agonist | GPCR | HEK293 | Kidney | Agonist | Fluorescence |
| tox21-kiss1r-wt-p2 | KISS1R wild type | GPCR | HEK293 | Kidney | Agonist | Fluorescence |
| tox21-ks-are-p1 | Nrf2/ARE | SR | HaCaT | Skin | Agonist | Luciferase |
| tox21-mitotox-p1 | Mitochondria toxicity | SR | HepG2 | Liver | Inhibition | Fluorescence |
| tox21-ms-ache-p2 | AChE (with human microsomes) | Neurotoxicity | NA | Biochemical | Inhibition | Colorimetric |
| tox21-ms-p53-p1 | P53 (with rat microsomes) | Gene Tox | HCT-116 | Colon Cancer | Agonist | Beta-lactamase |
| tox21-ms-p53-p2 | P53 (with human microsomes) | Gene Tox | HCT-116 | Colon Cancer | Agonist | Beta-lactamase |
| tox21-nfkb-bla-agonist-p1 | NFkB agonist | SR | ME-180 | Cervical Cancer | Agonist | Beta-lactamase |
| tox21-p450-1a2-p1 | CYP1A2 | Metabolism | NA | Biochemical | Inhibition | Luminescence |
| tox21-p450-2c19-p1 | CYP2C19 | Metabolism | NA | Biochemical | Inhibition | Luminescence |
| tox21-p450-2c9-p1 | CYP2C9 | Metabolism | NA | Biochemical | Inhibition | Luminescence |
| tox21-p450-2d6-p1 | CYP2D6 | Metabolism | NA | Biochemical | Inhibition | Luminescence |
| tox21-p450-3a4-p1 | CYP3A4 | Metabolism | NA | Biochemical | Inhibition | Luminescence |
| tox21-p53-bla-p1 | P53 | Gene Tox | HCT-116 | Colon Cancer | Agonist | Beta-lactamase |
| tox21-p53-bla-p7 | P53 | Gene Tox | HCT-116 | Colon Cancer | Agonist | Beta-lactamase |
| tox21-pgc-err-p1 | PGC-ERR | NR | HEK293 | Kidney | Agonist | Luminescence |

| | | | | | | |
|---|---|---|---|---|---|---|
| tox21-ppard-bla-agonist-p1 | PPAR-delta-BLA agonist | NR | HEK293 | Kidney | Agonist | Beta-lactamase |
| tox21-ppard-bla-antagonist-p1 | PPAR-delta-BLA antagonist | NR | HEK293 | Kidney | Antagonist | Beta-lactamase |
| tox21-pparg-bla-agonist-p1 | PPAR-gamma agonist | NR | HEK293 | Kidney | Agonist | Beta-lactamase |
| tox21-pparg-bla-antagonist-p1 | PPAR-gamma antagonist | NR | HEK293 | Kidney | Antagonist | Beta-lactamase |
| tox21-pr-bla-agonist-p1 | PR-BLA agonist | NR | HEK293 | Kidney | Agonist | Beta-lactamase |
| tox21-pr-bla-antagonist-p1 | PR-BLA antagonist | NR | HEK293 | Kidney | Antagonist | Beta-lactamase |
| tox21-pxr-p1 | PXR agonist | NR | HepG2 | Liver | Agonist | Luciferase |
| tox21-rar-agonist-p1 | RAR agonist | NR | C3H10T1/2 | Murine embryo fibroblast | Agonist | Luciferase |
| tox21-rar-antagonist-p2 | RAR antagonist | NR | C3H10T1/2 | Murine embryo fibroblast | Antagonist | Luciferase |
| tox21-rar-viability-p2 | RAR viability | Cytotoxicity | C3H10T1/2 | Murine embryo fibroblast | Cytotoxicity | Luciferase |
| tox21-ror-cho-antagonist-p1 | ROR antagonist | NR | CHO | Hamster | Antagonist | Luciferase |
| tox21-ror-cho-viability-p1 | ROR viability | Cytotoxicity | CHO | Hamster | Cytotoxicity | Fluorescence |
| tox21-rt-viability-hek293-p1-flor_8h_n | Cell viability | Cytotoxicity | HEK293 | Kidney | Cytotoxicity | Fluorescence |
| tox21-rt-viability-hek293-p1-flor_16h_n | Cell viability | Cytotoxicity | HEK293 | Kidney | Cytotoxicity | Fluorescence |
| tox21-rt-viability-hek293-p1-flor_24h_n | Cell viability | Cytotoxicity | HEK293 | Kidney | Cytotoxicity | Fluorescence |
| tox21-rt-viability-hek293-p1-glo_8h_n | Cell viability | Cytotoxicity | HEK293 | Kidney | Cytotoxicity | Luminescence |
| tox21-rt-viability-hek293-p1-glo_16h_n | Cell viability | Cytotoxicity | HEK293 | Kidney | Cytotoxicity | Luminescence |
| tox21-rt-viability-hek293-p1-glo_24h_n | Cell viability | Cytotoxicity | HEK293 | Kidney | Cytotoxicity | Luminescence |
| tox21-rt-viability-hepg2-p1-flor_8h_n | Cell viability | Cytotoxicity | HepG2 | Liver | Cytotoxicity | Fluorescence |
| tox21-rt-viability-hepg2-p1-flor_16h_n | Cell viability | Cytotoxicity | HepG2 | Liver | Cytotoxicity | Fluorescence |
| tox21-rt-viability-hepg2-p1-flor_24h_n | Cell viability | Cytotoxicity | HepG2 | Liver | Cytotoxicity | Fluorescence |
| tox21-rt-viability-hepg2-p1-glo_8h_n | Cell viability | Cytotoxicity | HepG2 | Liver | Cytotoxicity | Luminescence |
| tox21-rt-viability-hepg2-p1-glo_16h_n | Cell viability | Cytotoxicity | HepG2 | Liver | Cytotoxicity | Luminescence |
| tox21-rt-viability-hepg2-p1-glo_24h_n | Cell viability | Cytotoxicity | HepG2 | Liver | Cytotoxicity | Luminescence |
| tox21-rxr-bla-agonist-p1 | RXR-BLA | NR | HEK293 | Kidney | Agonist | Beta-lactamase |

| | | | | | | |
|---|---|---|---|---|---|---|
| tox21-sbe-bla-agonist-p1 | SBE-BLA (TGF-beta) agonist | Developmental Toxicity | HEK293 | Kidney | Agonist | Beta-lactamase |
| tox21-sbe-bla-antagonist-p1 | SBE-BLA (TGF-beta) antagonist | Developmental Toxicity | HEK293 | Kidney | Antagonist | Beta-lactamase |
| tox21-shh-3t3-gli3-agonist-p1 | Hedgehog agonist | Developmental Toxicity | NIH/3T3 | Murine embryo fibroblast | Agonist | Luciferase |
| tox21-shh-3t3-gli3-antagonist-p1 | Hedgehog antagonist | Developmental Toxicity | NIH/3T3 | Murine embryo fibroblast | Antagonist | Luciferase |
| tox21-spec-hek293-p1 | Auto Fluorescence | Counter Screen | HEK293 | Kidney | Counter | Fluorescence |
| tox21-spec-hepg2-p1 | Auto Fluorescence | Counter Screen | HepG2 | Liver | Counter | Fluorescence |
| tox21-trhr-hek293-p1 | TRHR agonist and antagonist | GPCR | HEK293 | Kidney | Agonist and Antagonist | Fluorescence |
| tox21-tshr-agonist-p1 | TSHR agonist | GPCR | HEK293 | Kidney | Agonist | Fluorescence |
| tox21-tshr-antagonist-p1 | TSHR antagonist | GPCR | HEK293 | Kidney | Antagonist | Fluorescence |
| tox21-tshr-wt-p1 | TSHR wild type | GPCR | HEK293 | Kidney | Agonist | Fluorescence |
| tox21-vdr-bla-agonist-p1 | VDR-BLA agonist | NR | HEK293 | Kidney | Agonist | Beta-lactamase |
| tox21-vdr-bla-antagonist-p1 | VDR-BLA antagonist | NR | HEK293 | Kidney | Antagonist | Beta-lactamase |

**Tox21 Data Curation and Filtering**

We began with 7,947,446 Tox21 concentration–response curves and applied a series of structural, data-quality, and assay-content filters. First, we removed curves associated with very small chemical structures (10,506 with fewer than three heavy atoms), ambiguous concentration annotations [3,073 lacking a `curve_class2(cc2)` label or with `cc2` > 4], and low-purity samples (2,026,038 with purity other than A or B). We then filtered on assay content encoded in SAMPLE_DATA_TYPE, excluding raw channel readouts (2,770,160), viability readouts (1,748,332), and FITC- or rhodamine-labeled channels (35,328 each). These endpoints were excluded because they are either handled differently in downstream analyses—for example, ratio metrics derived from paired channels such as ch1 and ch2, or used as counterscreens, such as viability measurements in mechanistic agonist/antagonist classification. After all filtering steps, 2,499,594 curves remained for the downstream analysis.

**Data Quality Control**
We applied the following quality-control filters to ensure data reliability:

1. Valid-response threshold: each curve was required to have at least twelve valid (non-missing) response measurements out of 15 concentration points.

2. Consistency enforcement: for curves classified as `Curve_class2(cc2)=4` (flat/inactive), we enforced consistency by setting Assay_outcome to "inactive" and $AC_{50}$ to NaN. Rows where `cc2=4` and the recorded Assay_outcome were inconsistent were demoted to inconclusive.

3. Sign-consistency check: agonist-labelled replicates with negative cc2 and antagonist-labelled replicates with positive `cc2` (994 and 1,023 rows respectively) were demoted to inconclusive with $AC_{50}$ set to NaN.

**Branch Structure**
The 102 assays were partitioned into mechanistic and non-mechanistic branches according to assay mode (Table S1). This branch-level formulation was designed to enable assay-mode-specific prediction consistent with the biological interpretation of the underlying readout. Mechanistic assays were modeled as a three-class classification task with labels agonist, antagonist, and inactive, whereas non-mechanistic assays were treated as a binary classification task with labels active and inactive. Mechanistic-branch outcomes containing the term "cytotoxic" were excluded because potential cytotoxicity could interfere with the primary mechanistic response, and such responses can resemble antagonist-like profiles. Inconclusive agonists were merged into the agonist class and inconclusive antagonists into the antagonist class, whereas records labeled only as inconclusive were excluded.

**Outcome-label collapsing**

The eight original assay outcome strings ("active agonist", "active antagonist", "inactive", "inconclusive", "inconclusive agonist", "inconclusive agonist (cytotoxic)", "inconclusive antagonist", "inconclusive antagonist (cytotoxic)" recorded in Tox21 records are mapped to five collapsed classes (Table S2).

For the mechanism branch, only the three primary classes - agonist, antagonist, and inactive - are used as supervised outcome labels during both pretraining and probing. The cytotoxic and inconclusive classes are excluded from outcome and $AC_{50}$ supervision, but their rows are retained for the masked-response reconstruction objective. For the non-mechanism branch, only the two primary classes - active and inactive - are used as supervised outcome labels during both pretraining and probing.

**Table S2. Outcome-label collapsing strategy**

| Original Tox21 outcome string | Collapsed class | Supervised? | Notes |
|---|---|---|---|
| `active agonist`<br>`inconclusive agonist` | Agonist | Yes | Both rows treated as the active-agonist class for outcome supervision. |
| `active antagonist`<br>`inconclusive antagonist` | Antagonist | Yes | Both rows treated as the active-antagonist class for outcome supervision. |
| `inactive` | Inactive | Yes | Used as the inactive class for outcome supervision. |
| `inconclusive agonist (cytotoxic)`<br>`inconclusive antagonist (cytotoxic)` | Cytotoxic | No | Excluded from outcome and $AC_{50}$ supervision; cytotoxicity confounds the receptor signal. |
| `inconclusive` | Inconclusive | No | Excluded from outcome and $AC_{50}$ supervision; retained for masked-response pretraining only. |

**Target Variables**

We prepared three complementary prediction targets: (1) concentration response values: normalized continuous responses at each concentration point for masked-response prediction; (2) assay outcome classification: agonist, antagonist, and inactive for the mechanism branch and active and inactive for the non-mechanism branch; (3) $AC_{50}$ regression, $\log_{10}$-transformed $AC_{50}$ values for compounds with active responses (zero $AC_{50}$ values are treated as NaN).

**Model architecture**

**Categorical embeddings**

Six assay descriptors: `Protocol_name`, `Target_category`, `Cell_type`, `Mode`, `Technology`, and `is_ratio`, each use a learned embedding table. In our implementation, `Protocol_name` uses embedding width 64 and the other five use width 32; each table is linearly projected to the model hidden size 768, and the six projected vectors are summed to form a single assay-context vector `assay_ctx`.

**Dose–response token encoding**

For each of the 15 concentration positions we build a token as follows:

**Dose** (**concentration**): $\log_{10}$ concentration is passed through a two-layer `MLP` (1 → 256 → 768) with `ReLU` between layers. Response: the per-dose normalized response is passed through a separate two-layer `MLP` (1 → 256 → 768) with `ReLU`.

**Token**: dose (concentration) and response embeddings are added to `assay_ctx` (broadcast along the dose axis) and passed through LayerNorm:

`token_i=LayerNorm(assay_ctx+dose_emb_i+response_emb_i)`

(An ablation can omit `assay_ctx` from DR tokens; the default includes it.)

**Special tokens**

A learnable [`CLS`] vector is added to `assay_ctx` and LayerNorm-normalized: [`CLS`] = LayerNorm(`CLS_base` + `assay_ctx`). At masked pretraining positions, the response channel is replaced by a learnable [`MASK`] embedding before the response `MLP` output is combined into the token.

**Transformer encoder**

The 16-token sequence ([`CLS`] plus 15 dose–response tokens) is encoded by a `TransformerEncoder`. In the released configuration we use `ASSAY_NUM_LAYERS = 6` encoder layers (six stacked `TransformerEncoderLayer` blocks), eight attention heads, hidden size 768, feed-forward size 4 × 768 = 3,072, dropout 0.1, pre-norm (`norm_first=True`), and `GELU` activation inside the encoder blocks.

Masked response prediction: a linear 768 → 1 head applied to each DR (dose response) token. Outcome classification: branch-specific linear heads applied during Phase 1 when auxiliary supervision is enabled ($\lambda = 0.1$). For the mechanism branch, a linear 768 → 3 head predicts inactive, agonist, or antagonist. For the non-mechanism branch, a linear 768 → 2 head predicts inactive or active. Cytotoxic and inconclusive rows are excluded from the supervised outcome loss.

**Branch probe heads (evaluation only)**

After Phase 1, we attach two linear probe heads on the frozen `CLS` vector: a 3-class mechanism head (agonist, antagonist, inactive) and a 2-class non-mechanism head (active, inactive). These are for evaluation, not Phase 1 optimization.

**Training procedure**

**Phase 1: backbone pre-training**

Each step randomly masks 20% of valid response positions; masked sites use the [`MASK`] embedding. The model is trained to reconstruct normalized responses at masked positions (masked MSE). Unknown / inconclusive outcomes (label −1) are excluded from supervised terms; supervised terms also restrict to rows whose cc2 lies in the allowed replicate-aggregation whitelist for the chosen `use_cc2` mode (strict vs loose). Under `cc2=strict`, the allowed set is (±1.1, ±1.2, ±2.1, ±2.2, ±4.0); loose adds additional cc2 classes, i.e., ±1.3, ±1.4, ±2.3, ±2.4, ±3.0, which is the full valid cc2 list.

**Optimization:** AdamW, learning rate $1 \times 10^{-4}$, weight decay $1 \times 10^{-4}$, 50 epochs. Default batch size is 4,096.

**Checkpoint selection**

For all pretraining variants, the backbone checkpoint was selected based on the lowest validation masked-response MSE. Validation outcome accuracy was recorded for monitoring purposes but was not used for checkpoint selection. This ensured that model selection remained consistent across variants and was tied to the shared reconstruction objective.

**Phase 2: outcome classification probe**

We load assay_backbone_best.pt, freeze the backbone, and train only the two probe heads for 5 epochs (AdamW, LR $1 \times 10^{-3}$, weight decay $1 \times 10^{-4}$). Mechanism branch: focal loss with $\gamma = 2.0$ and class $\alpha$ vector [1, 1, 1] for (agonist, antagonist, inactive). Non-mechanism branch: cross-entropy. Checkpoint selection: maximize validation 0.5·(macro-F1_mechanism + binary-F1_non-mechanism).

**Phase 3: $AC_{50}$ regression probe**

Using the same backbone checkpoint selected by validation masked-response MSE, we froze the backbone and trained the built-in CLS AC50 head (head_ac50) for 5 epochs using AdamW with a learning rate of $10^{-3}$. The training criterion is mean squared error on $\log_{10} AC_{50}$ for rows with finite ac50_log (raw $AC_{50} > 0$ before the log).

**Phase 4: $AC_{50}$ probe evaluation**

$AC_{50}$ probe predictions were evaluated per-protocol and overall on the validation and test splits using RMSE, MAE, $R^2$, and Pearson correlation. Sample-weighted aggregates across the 102 protocols were additionally computed.

**Phase 5: foundation model**

For deployment, we retrained Tox21mer on the full quality-controlled dataset, including all compounds from the original training, validation, and test splits, using the same architecture and hyperparameters as in the split-based pretraining stage. The resulting full-data checkpoint was designated as the final foundation model for downstream applications. Because this retraining step used the entire dataset, no held-out set remained for model selection at this stage; accordingly, unbiased estimates of generalization remain those obtained from the original split-based evaluation. From the final full-data model, we extracted a 768-dimensional [`CLS`] embedding for each individual curve and then averaged embeddings across technical replicates for each unique (chemical, protocol) pair to generate the representations used in downstream analyses.

**Implementation details & computing**

All models were implemented in PyTorch 2.x. Random seeds were fixed (seed= 42 ) for all stochastic operations, and deterministic CUDA behavior was enabled by setting `torch.backends.cudnn.deterministic = True` and `torch.backends.cudnn.benchmark = False`. DataLoader workers were seeded using a `worker_init_fn` aligned with `torch.initial_seed()`. For full bitwise reproducibility on GPU, the environment variable `CUBLAS_WORKSPACE_CONFIG=:4096:8` was set. Mixed-precision training was used where applicable, and 8–16 DataLoader workers were employed to improve throughput across multiple GPUs.

All training runs were performed on NVIDIA A100-SXM4 GPUs with either 40 GB or 80 GB of memory. A single Phase 1 training run (50 epochs; 1,736,846 training rows; batch size 4.096) required approximately 6 hours. Phase 5 retraining on the full dataset required an additional approximately 7 hours. The probe stages (Phases 2–4) together required approximately 15 minutes on a single GPU. Each `MLP` baseline completed in less than 2 hours.

**Additional Results**

**Results for `CC2=Loose`**

As mentioned before, we defined two cc2 groupings: `cc2 strict` and `cc2 loose`. The `cc2 strict` group comprised the original Tox21 classes (-1.1, -1.2, -2.1, -2.2, 1.1, 1.2, 2.1, 2.2, and 4), whereas the cc2 loose group included 1.1, 1.2, 1.3, 1.4, 2.1, 2.2, 2.3, 2.4, 3.0, 4.0, -1.1,-1.2, -1.3, -1.4, -2.1, -2.2, -2.3, -2.4. In the main manuscript, all probing analyses and model training were conducted using `cc2=strict`. Below, we provide summary results for `cc2=loose`.

We compared branch-classification performance under `cc2=strict` (**Table 3a**, **3b**) versus `cc2= loose` (**Table S3a** and **3b**). Overall, the strict setting performed substantially better: the composite score increased from 0.9607 under `cc2 =loose` to 0.9891 under cc2=strict, a difference of 2.84 points. Most of this gap was driven by the mechanism branch, where macro-F1 declined from 0.9848 to 0.9467 when moving from strict to loose. By comparison, the non-mechanism branch showed a smaller decrease, with binary F1 falling from 0.9935 to 0.9747. Test-set support also changed, particularly for mechanism-active classes—because the loose definition retains additional cc2 patterns excluded under the strict definition (e.g., agonist/antagonist support increased from 9,302 / 7,180 under strict to 12,158 / 10,305 under loose). Thus, part of the difference reflects the broader and more challenging label space under cc2 = loose, rather than a change in model performance on an identical set of rows.

Among mechanism classes, agonist prediction was affected most strongly by the loose setting. Agonist F1 decreased from 0.9779 to 0.8988, driven primarily by a drop in precision from 0.9810 to 0.8251, while recall remained high (0.9750 to 0.9868). This pattern is consistent with increased overcalling of inactive compounds as agonists. For example, the confusion matrix shows 144,674 correct inactive predictions under loose versus 147,426 under strict, with 2,506 + 566 inactive rows misclassified as agonist or antagonist under loose compared with only 162 + 158 under strict. Antagonist performance also worsened under the loose setting, with F1 declining from 0.9787 to 0.9535.

The non-mechanism branch showed the same general trend, although at a smaller scale. Accuracy declined from 0.9899 under strict to 0.9643 under loose, accompanied by more active/inactive confusion: 778 active→inactive and 2,503 inactive→active errors under loose, compared with 218 and 638, respectively, under strict. Although the dominant error type differed, the total error burden was clearly larger under the loose condition.

Ranking metrics were consistent with these classification results. Per-class PR-AUC and ROC-AUC remained high in both settings but were uniformly better under cc2=strict (e.g., non-mechanism active PR-AUC: 0.9981 under strict vs. 0.9913 under loose). These results suggest that expanding the allowable cc2 set introduces cases that are less cleanly separable in feature space, rather than merely affecting metric reporting. Overall, cc2 =loose provides broader curve-type coverage but at the cost of noticeably reduced branch-classification performance, with the mechanism branch and especially agonist versus inactive discrimination showing the greatest degradation.

**Table S3a**. Test data mechanism and non-mechanism branch classification reports for cc2=loose.

| | **Mechanism branch** | | | |
|---|---|---|---|---|
| | Precision | Recall | F1-score | Support |
| Agonist | 0.8251 | 0.9868 | 0.8988 | 12,158 |
| Antagonist | 0.9445 | 0.9627 | 0.9535 | 10,305 |
| Inactive | 0.9966 | 0.9792 | 0.9878 | 147,746 |

| | | | | |
|---|---|---|---|---|
| Accuracy | | | 0.9958 | 170,209 |
| Macro Avg | 0.9221 | 0.9763 | 0.9848 | 170,209 |
| Weighted Avg | 0.9812 | 0.9788 | 0.9958 | 170,209 |
| | **Non-mechanism branch** | | | |
| Active | 0.9102 | 0.9703 | 0.9393 | 26,163 |
| Inactive | 0.9878 | 0.9619 | 0.9747 | 65,651 |
| Accuracy | | | 0.9899 | 91,814 |
| Macro Avg | 0.9490 | 0.9661 | 0.9570 | 91,814 |
| Weighted Avg | 0.9657 | 0.9643 | 0.9646 | 91,814 |

**Table S3b.** Per-class PR-AUC and ROC-AUC on the held-out test set for `cc2 =loose`.

| | Branch Class | PR-AUC | ROC-AUC | Support |
|---|---|---|---|---|
| mechanism | Agonist | 0.9899 | 0.9989 | 12,158 |
| | Antagonist | 0.9922 | 0.9993 | 10,305 |
| | Inactive | 0.9997 | 0.9981 | 147,746 |
| Non-mechanism | Active | 0.9913 | 0.9959 | 26,163 |

We also retrained $AC_{50}$ supervision for all curves with `cc2=loose` using an inverse weighted MSE loss as for the `cc2=strict` curves.

In the weighted $AC_{50}$ experiment, the CC2-restricted setting (`cc2=strict`) outperformed `cc2=loose` across both global (**Table S4**) and tail-focused metrics once CC2 gating was properly enforced during $AC_{50}$ training and evaluation (**Figure S2**). Under the strict setting, test performance was consistently stronger across all core regression metrics: $R^2$ = 0.8738 versus 0.7659, Pearson = 0.9423 versus 0.8981, MAE = 0.2011 versus 0.2912, and RMSE = 0.2894 versus 0.4088. Validation performance also improved substantially, with the best validation MSE decreasing from 0.1632 under loose to 0.0800 under strict.

Importantly, this comparison reflects different row pools by design (n = 52,417 for `cc2=strict` versus 83,143 for `cc2=loose`), because the strict setting excludes Curve_class2 patterns that are retained under the loose definition. Despite using the smaller and more constrained dataset, the strict model showed better calibration in the low-$AC_{50}$ regime, with lower low-tail MAE (0.733 versus 1.034) and a smaller low-tail residual mean (0.733 versus 1.004), indicating reduced positive bias for highly potent compounds. Overall, these weighted results support the conclusion that, for $AC_{50}$ regression, the stricter `cc2` definition provides a cleaner supervision subset and yields materially better predictive performance than the broader loose setting.

**Table S4**. Comparison in $AC_{50}$ supervision.

| Category | N | $R^2$ | Pearson | mae | rmse | Low tail mae |
|---|---|---|---|---|---|---|
| `cc2=strict` | 52,417 | 0.87 | 0.94 | 0.201 | 0.289 | 0.257 |
| `cc2=loose` | 83,143 | 0.77 | 0.90 | 0.291 | 0.409 | 1.034 |

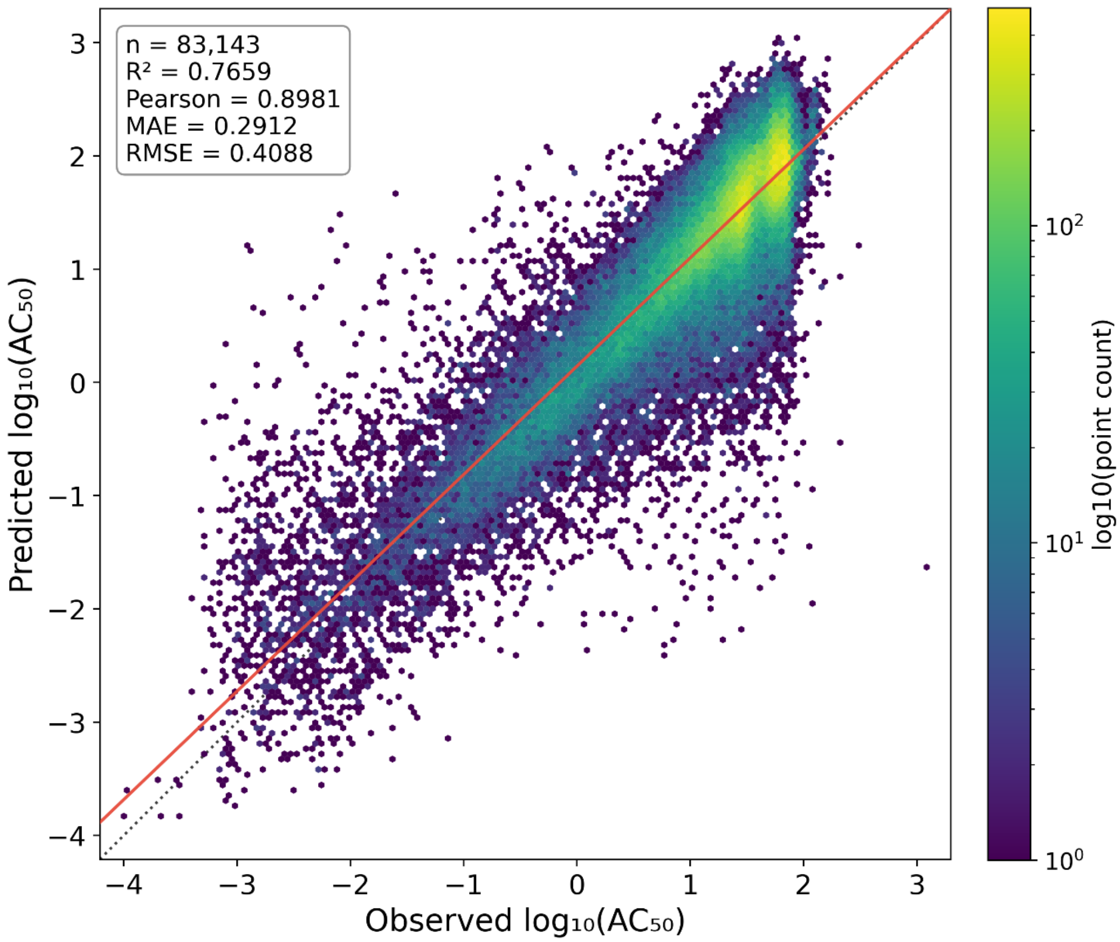


**Figure S2. Predicted log$_{10}$(AC$_{50}$) vs observed log$_{10}$(AC$_{50}$) values for all test curves with valid AC$_{50}$ values and `cc2=loose`** (Pearson and Spearman ρ = ~0.89).

As expected, the separation among the different `cc2` classes was less well defined than in the `cc2=loose` setting (**Figure S2**).

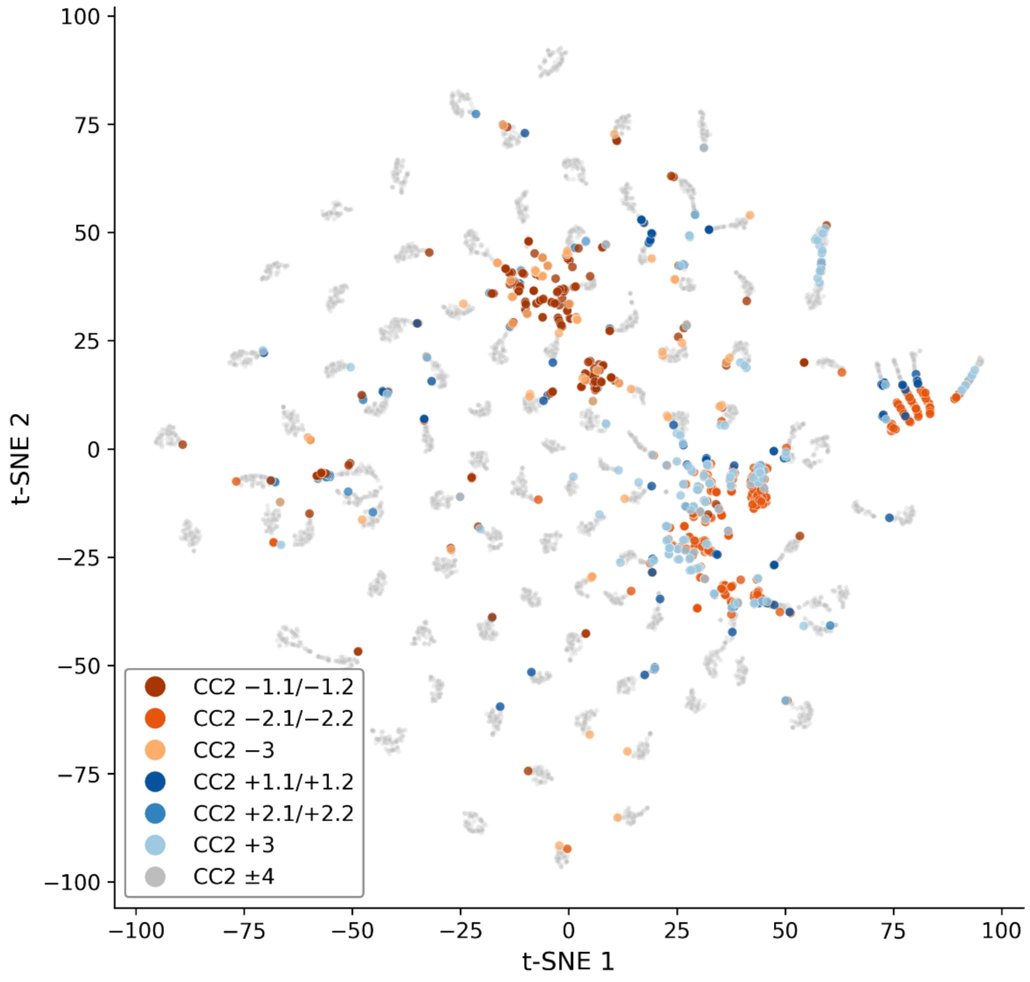


**Figure S3**. Two-dimensional t-SNE projection of the 768-dimensional embeddings for test curves shown for a stratified subsample of 8,000 of the 394,983 unique (SMILES, protocol) embeddings. The full set of curve class 2 (cc2) labels represented in the model was (1.1, 1.2, 1.3, 1.4, 2.1, 2.2, 2.3, 2.4, 3.0, -1.1, -1.2, -1.3, -1.4, -2.1, -2.2, -2.3, -2.4, -3.0, 4.0). For display purposes, these labels were consolidated into seven groups, shown in the legend.

**Table S4**. $AC_{50}$ Performance Metrics on Test Curves Using Inverse-Weighted MSE Loss During Training.

| Protocol name | n_samples | rmse | mae | $R^2$ | Pearson ρ |
|---|---|---|---|---|---|
| tox21-pr-bla-agonist-p1 | 143 | 0.2060 | 0.1611 | 0.9763 | 0.9898 |
| tox21-er-luc-bg1-4e2-agonist-p4 | 177 | 0.2224 | 0.1676 | 0.9720 | 0.9867 |
| tox21-ar-mda-kb2-luc-agonist-p3 | 252 | 0.2377 | 0.1873 | 0.9693 | 0.9847 |
| tox21-er-bla-agonist-p2 | 297 | 0.2383 | 0.1823 | 0.9655 | 0.9838 |
| tox21-gr-hela-bla-agonist-p1 | 315 | 0.2671 | 0.1825 | 0.9551 | 0.9775 |
| tox21-dt40-p1-657 | 1289 | 0.2043 | 0.1586 | 0.9511 | 0.9861 |
| tox21-erb-bla-p1 | 152 | 0.3057 | 0.2285 | 0.9497 | 0.9793 |
| tox21-dt40-p1-653 | 1309 | 0.2069 | 0.1585 | 0.9472 | 0.9845 |

| | | | | | |
|---|---|---|---|---|---|
| tox21-ar-mda-kb2-luc-agonist-p1 | 190 | 0.3440 | 0.2701 | 0.9404 | 0.9776 |
| tox21-p450-2c19-p1 | 1756 | 0.1692 | 0.1121 | 0.9358 | 0.9758 |
| tox21-drd2-antagonist-p1 | 171 | 0.2973 | 0.1919 | 0.9351 | 0.9704 |
| tox21-fxr-bla-antagonist-p1 | 480 | 0.2231 | 0.1438 | 0.9328 | 0.9672 |
| tox21-dt40-p1-100 | 1293 | 0.2303 | 0.2008 | 0.9316 | 0.9901 |
| tox21-pr-bla-antagonist-p1 | 1229 | 0.1692 | 0.1354 | 0.9305 | 0.9798 |
| tox21-aromatase-p1 | 893 | 0.2773 | 0.2076 | 0.9299 | 0.9738 |
| tox21-p450-3a4-p1 | 1451 | 0.1815 | 0.1391 | 0.9269 | 0.9709 |
| tox21-er-luc-bg1-4e2-agonist-p2 | 806 | 0.2716 | 0.1870 | 0.9252 | 0.9660 |
| tox21-ar-mda-kb2-luc-antagonist-p1 | 780 | 0.3126 | 0.2186 | 0.9179 | 0.9641 |
| tox21-ror-cho-antagonist-p1 | 828 | 0.2179 | 0.1574 | 0.9173 | 0.9647 |
| tox21-rxr-bla-agonist-p1 | 521 | 0.2316 | 0.1706 | 0.9145 | 0.9579 |
| tox21-rar-agonist-p1 | 386 | 0.2878 | 0.2178 | 0.9124 | 0.9567 |
| tox21-casp3-cho-p1 | 130 | 0.2393 | 0.1879 | 0.9114 | 0.9623 |
| tox21-p450-2d6-p1 | 1435 | 0.2129 | 0.1735 | 0.9058 | 0.9822 |
| tox21-shh-3t3-gli3-antagonist-p1 | 1139 | 0.2435 | 0.1772 | 0.9033 | 0.9659 |
| tox21-casp3-hepg2-p1 | 327 | 0.2278 | 0.1638 | 0.9010 | 0.9538 |
| tox21-ar-bla-agonist-p1 | 309 | 0.4131 | 0.3188 | 0.8986 | 0.9613 |
| tox21-sbe-bla-antagonist-p1 | 474 | 0.2597 | 0.2174 | 0.8952 | 0.9820 |
| tox21-p450-1a2-p1 | 2174 | 0.1634 | 0.1358 | 0.8944 | 0.9788 |
| tox21-car-antagonist-p1 | 1297 | 0.2241 | 0.1674 | 0.8910 | 0.9533 |
| tox21-tshr-wt-p1 | 38 | 0.5135 | 0.4238 | 0.8864 | 0.9729 |
| tox21-ks-are-p1 | 574 | 0.2861 | 0.2073 | 0.8818 | 0.9502 |
| tox21-ar-mda-kb2-luc-antagonist-p2 | 1211 | 0.2903 | 0.1772 | 0.8798 | 0.9551 |
| tox21-ache-p4 | 94 | 0.2290 | 0.1751 | 0.8785 | 0.9549 |
| tox21-rt-viability-hek293-p1-glo_16h_n | 252 | 0.2142 | 0.1431 | 0.8757 | 0.9450 |
| tox21-pparg-bla-antagonist-p1 | 501 | 0.2570 | 0.1902 | 0.8746 | 0.9381 |
| tox21-mitotox-p1 | 1115 | 0.2385 | 0.1488 | 0.8642 | 0.9336 |
| tox21-ache-p5 | 62 | 0.2384 | 0.1845 | 0.8641 | 0.9554 |
| tox21-er-luc-bg1-4e2-antagonist-p2 | 827 | 0.2805 | 0.1978 | 0.8612 | 0.9477 |
| tox21-err-p1 | 2406 | 0.2505 | 0.1632 | 0.8585 | 0.9332 |
| tox21-er-luc-bg1-4e2-antagonist-p1 | 631 | 0.3415 | 0.2446 | 0.8573 | 0.9434 |
| tox21-rar-antagonist-p2 | 1021 | 0.2845 | 0.2148 | 0.8552 | 0.9482 |
| tox21-gh3-tre-antagonist-p1 | 1008 | 0.2765 | 0.2285 | 0.8506 | 0.9696 |
| tox21-cre-agonist-p1 | 39 | 0.4506 | 0.3338 | 0.8499 | 0.9346 |
| tox21-gr-hela-bla-antagonist-p1 | 635 | 0.3784 | 0.2633 | 0.8487 | 0.9221 |
| tox21-h2ax-cho-p2 | 327 | 0.2816 | 0.1766 | 0.8463 | 0.9278 |
| tox21-ppard-bla-antagonist-p1 | 291 | 0.3627 | 0.2336 | 0.8389 | 0.9366 |
| tox21-erb-bla-antagonist-p1 | 865 | 0.2738 | 0.1455 | 0.8377 | 0.9171 |
| tox21-chrm1-p1 | 293 | 0.4185 | 0.2928 | 0.8353 | 0.9216 |
| tox21-tshr-agonist-p1 | 273 | 0.2904 | 0.2124 | 0.8346 | 0.9256 |
| tox21-herg-u2os-p1 | 576 | 0.2328 | 0.1908 | 0.8285 | 0.9693 |
| tox21-ahr-p1 | 588 | 0.2566 | 0.1877 | 0.8248 | 0.9248 |
| tox21-pxr-p1 | 1347 | 0.2445 | 0.1638 | 0.8244 | 0.9190 |
| tox21-esre-bla-p1 | 228 | 0.4998 | 0.3604 | 0.8227 | 0.9084 |
| tox21-rt-viability-hek293-p1-glo_8h_n | 181 | 0.1675 | 0.1246 | 0.8205 | 0.9499 |
| tox21-htr2a-p1 | 654 | 0.4513 | 0.3455 | 0.8199 | 0.9491 |
| tox21-pparg-bla-agonist-p1 | 410 | 0.3413 | 0.2544 | 0.8036 | 0.9006 |

| | | | | | |
|---|---|---|---|---|---|
| tox21-p450-2c9-p1 | 1892 | 0.2242 | 0.1730 | 0.8014 | 0.9421 |
| tox21-ap1-agonist-p1 | 427 | 0.2771 | 0.2224 | 0.7961 | 0.9416 |
| tox21-are-bla-p1 | 922 | 0.2874 | 0.2256 | 0.7956 | 0.9286 |
| tox21-rt-viability-hek293-p1-glo_24h_n | 287 | 0.2955 | 0.1696 | 0.7932 | 0.9229 |
| tox21-car-agonist-p1 | 859 | 0.2641 | 0.2137 | 0.7810 | 0.9375 |
| tox21-p53-bla-p7 | 132 | 0.3727 | 0.2654 | 0.7799 | 0.9216 |
| tox21-cre-antagonist-p1 | 436 | 0.2625 | 0.1896 | 0.7777 | 0.9217 |
| tox21-ar-bla-antagonist-p1 | 905 | 0.3579 | 0.2486 | 0.7728 | 0.9328 |
| tox21-ache-p3 | 335 | 0.2996 | 0.2317 | 0.7713 | 0.9485 |
| tox21-ror-cho-viability-p1 | 438 | 0.3325 | 0.2177 | 0.7708 | 0.9085 |
| tox21-gnrhr-hek293-p2 | 195 | 0.5156 | 0.3874 | 0.7707 | 0.8791 |
| tox21-p53-bla-p1 | 461 | 0.4105 | 0.2574 | 0.7705 | 0.9002 |
| tox21-vdr-bla-antagonist-p1 | 329 | 0.4836 | 0.3272 | 0.7703 | 0.9093 |
| tox21-pgc-err-p1 | 1726 | 0.3383 | 0.2500 | 0.7635 | 0.8967 |
| tox21-elg1-luc-agonist-p1 | 115 | 0.4824 | 0.3729 | 0.7574 | 0.8809 |
| tox21-adrb2-antagonist-p1 | 148 | 0.5259 | 0.4153 | 0.7532 | 0.9298 |
| tox21-shh-3t3-gli3-agonist-p1 | 127 | 0.4317 | 0.3117 | 0.7389 | 0.8934 |
| tox21-rar-viability-p2 | 283 | 0.3487 | 0.2678 | 0.7305 | 0.8702 |
| tox21-hse-bla-p1 | 283 | 0.4272 | 0.2561 | 0.7240 | 0.8919 |
| tox21-ppard-bla-agonist-p1 | 128 | 0.4834 | 0.3775 | 0.7060 | 0.8440 |
| tox21-er-bla-antagonist-p1 | 544 | 0.4746 | 0.2909 | 0.6958 | 0.8774 |
| tox21-adrb2-agonist-p1 | 202 | 0.5563 | 0.4219 | 0.6955 | 0.8825 |
| tox21-ms-p53-p2 | 87 | 0.3323 | 0.2739 | 0.6951 | 0.9067 |
| tox21-gh3-tre-agonist-p1 | 39 | 0.6120 | 0.5315 | 0.6803 | 0.9522 |
| tox21-vdr-bla-agonist-p1 | 128 | 0.4931 | 0.4237 | 0.6637 | 0.8342 |
| tox21-kiss1r-hek293-p2 | 256 | 0.5978 | 0.4035 | 0.6341 | 0.8301 |
| tox21-rt-viability-hepg2-p1-glo_16h_n | 93 | 0.2609 | 0.1735 | 0.6032 | 0.8121 |
| tox21-drd2-agonist-p1 | 35 | 0.8114 | 0.6692 | 0.5982 | 0.9047 |
| tox21-nfkb-bla-agonist-p1 | 67 | 0.6302 | 0.4140 | 0.5302 | 0.7820 |
| tox21-hdac-p1 | 319 | 0.3306 | 0.2764 | 0.5301 | 0.9495 |
| tox21-rt-viability-hepg2-p1-flor_24h_n | 137 | 0.2545 | 0.1927 | 0.5251 | 0.8257 |
| tox21-rt-viability-hepg2-p1-glo_24h_n | 96 | 0.3387 | 0.2296 | 0.5231 | 0.8491 |
| tox21-hre-bla-agonist-p1 | 172 | 0.3445 | 0.2441 | 0.4973 | 0.8719 |
| tox21-rt-viability-hepg2-p1-flor_8h_n | 81 | 0.3329 | 0.2723 | 0.4626 | 0.7629 |
| tox21-rt-viability-hepg2-p1-glo_8h_n | 83 | 0.2701 | 0.1860 | 0.4279 | 0.7421 |
| tox21-ms-ache-p2 | 41 | 0.5098 | 0.3476 | 0.4257 | 0.8089 |
| tox21-ms-p53-p1 | 85 | 0.4509 | 0.3132 | 0.4234 | 0.8542 |
| tox21-rt-viability-hek293-p1-flor_8h_n | 51 | 0.2894 | 0.2449 | 0.4190 | 0.7638 |
| tox21-rt-viability-hepg2-p1-flor_16h_n | 111 | 0.3973 | 0.3322 | 0.4159 | 0.8436 |
| tox21-tshr-antagonist-p1 | 205 | 0.3777 | 0.2679 | 0.3868 | 0.7647 |
| tox21-rt-viability-hek293-p1-flor_16h_n | 80 | 0.3863 | 0.3344 | 0.3311 | 0.8140 |
| tox21-trhr-hek293-p1 | 359 | 0.4761 | 0.3349 | 0.2732 | 0.7032 |
| tox21-rt-viability-hek293-p1-flor_24h_n | 100 | 0.3703 | 0.3318 | 0.1484 | 0.8643 |
| tox21-kiss1r-wt-p2 | 54 | 0.2676 | 0.2056 | -0.1582 | 0.4470 |
| tox21-fxr-bla-agonist-p2 | 85 | 0.5483 | 0.3928 | -0.1857 | 0.6351 |
| tox21-sbe-bla-agonist-p1 | 29 | 0.5550 | 0.4585 | -5.7981 | 0.3796 |

**Self-supervised pretraining alone yields a strong representation**

Without auxiliary supervision, Tox21mer still performed strongly, reaching mechanism macro-F1 0.957 (agonist 0.925, antagonist 0.952, inactive 0.998) and non-mechanism binary-F1 0.966 (active 0.947, inactive 0.986). PR-AUC remained high for all classes, exceeding 0.977 throughout. Relative to the auxiliary-supervised model.

**Table S5.** Effect of auxiliary supervised losses on branch-probe performance without auxiliary supervision.

| | Mech F1 | | | | Non-mech F1 | | |
|---|---|---|---|---|---|---|---|
| **Configuration** | **Macro Avg** | **Agonist** | **Antagonist** | **Inactive** | **Binary** | **Active** | **inactive** |
| Tox21mer (with auxiliary supervision) | 0.9848 | 0.9779 | 0.9787 | 0.9978 | 0.9856 | 0.9778 | 0.9856 |
| Tox21mer (no auxiliary supervision) | 0.9568 | 0.9246 | 0.9522 | 0.9937 | 0.9664 | 0.9472 | 0.9855 |
| Difference (supervision − none) | +0.028 | +0.053 | +0.027 | +0.004 | +0.019 | +0.031 | +0.001 |

**Additional Ablation Intervention**

In the main text, we reported three blocks of ablations designed to decouple the contributions of response values, response ordering, and assay context to the backbone's CLS representation (**Table S6**). To assess the contribution of assay-context metadata to probe performance, we repeated the Phase 2a evaluation with the six categorical context features (protocol name, target category, cell type, assay mode, technology, and ratio flag) zeroed at every dose–response token during the frozen-backbone forward pass, so that the CLS embedding was derived solely from the response values and log-dose inputs. This ablation produced mechanism macro-F1 of 0.986 and non-mechanism binary-F1 of 0.994, essentially indistinguishable from the full-context baseline (0.985 and 0.986, respectively) (**Table S6**). Per-class PR-AUC remained at or above 0.997 on all four classes, and the confusion matrices showed no systematic shift in error pattern. The near-zero effect of removing assay context is consistent with the observation that, within each protocol, the response-value distribution alone is already highly discriminative for outcome classification, protocol identity is largely redundant once the model observes the response magnitudes. This result does not imply that assay context is uninformative in general; it indicates that, for the branch-probe task as posed, the response values carry sufficient signal. Context may become more important for cross-protocol transfer or for tasks where protocol identity is not implicitly recoverable from the response distribution.

**Table S6.** Single-factor ablation with zero-context inputs applied during Phase 1 pretraining.

| Ablation flag | Mech F1 | Non-mech F1 | Active recall | Agonist recall | Antagonist recall |
|---|---|---|---|---|---|
| None (baseline) | 0.9848 | 0.9935 | 0.9856 | 0.9750 | 0.9815 |
| `--zero-dr-assay-ctx` | 0.9857 | 0.9861 | 0.9877 | 0.9816 | 0.9850 |

**Table S7.** Transformer vs MLP baseline performance on the test set, using identical branch-probe supervision, label masking, and metrics. The 2-layer MLP on raw `resp_norm` plus `assay_ctx` matches the transformer to within one F1 point on both heads.

| Model | Features | Mech F1 | Non-mech F1 | Score |
|---|---|---|---|---|
| Tox21mer | Backbone CLS | 0.985 | 0.986 | 0.991 |
| MLP (stats only) | 8 summary stats | 0.846 | 0.965 | 0.906 |
| MLP (stats + ctx) | 8 stats + `assay_ctx` | 0.948 | 0.989 | 0.969 |
| MLP (raw only) | 15 Raw `resp_norm` | 0.910 | 0.981 | 0.945 |
| MLP (raw + ctx) | Raw `resp_norm` + `assay_ctx` | 0.983 | 0.996 | 0.990 |

**Additional Discussion on Ablations**

The contrast between Phases 2b and 2d indicates a marked asymmetry: evaluating a clean-trained probe on shuffled inputs reduced performance much more than evaluating a shuffled-trained probe on clean inputs. This pattern suggests that the observed drop under shuffling is driven mainly by probe-level distribution shift rather than by complete loss of predictive information in the backbone representation. Consistent with this interpretation, probes trained on shuffled curves recovered near-baseline performance, indicating that substantial outcome-relevant information is retained even when dose–response ordering is disrupted.

The MLP comparison supports a similar conclusion. Raw-response MLPs, especially when combined with assay context, achieved probe performance close to that of the transformer. Together, these results suggest that the probe tasks rely heavily on response-value distributions and assay context, while being less sensitive to fine-grained within-curve ordering. This does not diminish the broader utility of Tox21mer, but it does clarify the specific aspects of representation quality captured by the probe-based evaluation.

**Additional Discussion on Supervision**

We verified that probe-evaluation performance does not arise from circular reuse of pretraining supervision, we additionally pretrained Tox21mer with all auxiliary supervised loss weights set to zero (masked-response reconstruction only). The resulting purely self-supervised backbone retained mechanism macro-F1 of 0.957 and non-mechanism binary-F1 of 0.986, with per-class PR-AUC at or above 0.977 on every class. Auxiliary supervision contributes 1–6 F1 points of improvement, concentrated on the minority active classes, consistent with the interpretation that auxiliary terms sharpen minority-class decision boundaries beyond what reconstruction alone recovers. Probe metrics therefore measure compound-level generalization rather than retention of supervised pretraining signal. We therefore interpret the probe metrics as measuring generalization across held-out compounds, not simple memorization of the auxiliary task family. Second, pretraining objectives that explicitly reward shape-aware representations, for example, contrastive objectives that distinguish permutations of the same response bag, or auxiliary regression of fitted Hill-slope parameters, may produce embeddings that are more informative on tasks where shape is dispositive, such as mechanism-of-action classification across novel receptor classes. Third, adapting Tox21mer to related high-throughput assay resources, including ToxCast and recent concentration-response extensions of the Tox21 10K panel, would allow the representation to be exercised across a broader chemical and assay space and would clarify which aspects of the current model are dataset-specific and which generalize.
